# Modification-aware AI enables terminal chemical modifications for peptide design and discovers potent antimicrobials

**DOI:** 10.64898/2026.04.09.717597

**Authors:** Jing Xu, Marcelo D. T. Torres, Chen Li, Jian Li, Fuyi Li, Jiangning Song, Cesar de la Fuente-Nunez

## Abstract

Antimicrobial resistance (AMR) is an escalating global health threat, intensifying the urgent need for novel antibiotics. Here, we introduce Termini, a generative machine learning (ML) framework that integrates peptide generation, classification, and regression modules to systematically identify potent antimicrobial peptides (AMPs). A key feature of this pipeline is its explicit design of both N- and C-terminal modifications, combined with predictive modeling across 15 bacterial species. We experimentally validated the framework against 11 pathogenic species, representing the broadest spectrum of testing reported to date in such a study. We synthesized and tested 120 peptides (including 60 unique amino acid sequences) with and without terminal modifications. Remarkably, 111 of these peptides (92.5%) demonstrated antimicrobial activity *in vitro*. Comparative analyses demonstrated that terminal modifications substantially enhanced antimicrobial potency. Furthermore, the lead candidates identified through *in vitro* screening exhibited robust anti-infective efficacy in subsequent *in vivo* experiments, confirming their therapeutic potential. Collectively, these results highlight the high hit rate, biological relevance, and translational promise of this integrative, modification-aware AI framework for next-generation antimicrobial peptide development.

## Introduction

The rising threat of microbial pathogens, particularly drug-resistant strains, represents an increasingly serious and escalating challenge to global public health. Alarmingly, antimicrobial resistance (AMR) is projected to become the leading cause of death worldwide by 2050, surpassing fatalities from cancer and cardiovascular diseases ^1^. This crisis is largely driven by the overuse of antibiotics, which has accelerated the emergence and spread of resistant pathogens, thereby diminishing the arsenal of effective treatments^2^. Accordingly, there is an urgent need for new anti-infective agents with mechanisms that are less susceptible to resistance.

Antimicrobial peptides (AMPs) have emerged as promising alternatives to conventional antibiotics. Many AMPs exhibit broad-spectrum activity and act through multifaceted mechanisms, properties that may reduce the likelihood of resistance development ^3^. In parallel, advances in peptide synthesis and chemical modification now enable the rapid production and optimization of peptide candidates, supporting their evaluation as therapeutic agents ^4^.

AMPs typically carry high positive charges, with cationic amino acids forming a hydrophilic side and hydrophobic residues on the opposite side ^3,4^. Additionally, AMPs often possess alpha-helix and beta-sheet secondary structures ^5^. Together, these physicochemical properties facilitate interaction with and disruption of negatively charged bacterial membranes ^6^, a mode of action that can be difficult for bacteria to evade and therefore offers a potential route to counter multidrug-resistant infections ^6,7^.

Despite their promise, systematic AMPs discovery remains challenging because the design space is immense: AMPs are typically ≤50 amino acid residues long, and even restricting to the 20 canonical amino acids yields an astronomical number of possible sequences ^8^. To navigate this landscape, generative machine-learning approaches have been developed to propose candidate AMPs by learning patterns from known antimicrobial sequences, with the goal of accelerating discovery and improving development efficiency.

Several deep generative strategies have been explored for *de novo* AMP design. For instance, Das *et al.* proposed a variational autoencoder (VAE)-based framework coupled with a deep learning predictor to generate candidate peptides ^9^. Among 20 synthesized sequences, two exhibited antimicrobial activity against *Staphylococcus aureus* and *Escherichia coli*; however, their potency was modest, with HC_50_ values ranging from 125–500 µg mL^−1^ and LD_50_ values between 158–182 mg Kg^−1^. Szymczak *et al.* developed a conditional VAE (cVAE) to bias generation toward activity ^10^; of 33 synthesized peptides, 28 were active and 18 were non-toxic (HC_50_ ≥ 512 µg mL^−1^), underscoring the value of conditional generation for balancing efficacy and safety. In parallel, Cao *et al.* applied a generative adversarial network (GAN) integrated with BERT and multilayer perceptron (MLP) classifiers ^11^; although candidates were predicted to be active across multiple pathogens, only one of six synthesized peptides was active experimentally. More recently, Wang et al. introduced a diffusion-based generative model augmented with ensemble predictors (RNNs, attention, and CNNs) ^12^; of 40 synthesized candidates, 25 were active, and two lead peptides demonstrated broad-spectrum efficacy against bacterial and fungal pathogens, including *A. baumannii*, *K. pneumoniae*, *C. albicans*, and *C. glabrata*. Another recent study integrated ProGen2-xlarge with reinforcement learning fine-tuning to guide AMP design ^13^.

Although these approaches demonstrate the promise of generative models for AMP discovery, important limitations remain. Many pipelines do not explicitly model the effects of terminal modifications on antimicrobial potency, were validated against relatively narrow microbial panels, and often achieve modest success rates among synthesized candidates. To address these challenges, we developed an integrative, modification-aware framework for AMP discovery. Our approach used a diffusion-based generative model trained on an expanded AMP dataset to generate candidate sequences, followed by predictive screening for antimicrobial activity. Critically, we incorporated predictive models that explicitly account for N-terminal acetylation and C-terminal amidation during candidate evaluation, treating terminal chemistry as a first-class design variable rather than a *post hoc* optimization step. We rigorously validated the framework by synthesizing and testing 120 peptides *in vitro* and systematically quantifying how these common terminal modifications influence antimicrobial activity. By directly addressing these bottlenecks, our study establishes a scalable platform for identifying peptide-based therapeutics and expanding the urgently needed arsenal against multidrug-resistant pathogens.

## Results

### Generative modeling establishes a systematic pipeline for AMP discovery

The vast and diverse sequence space of potential AMPs presents a fundamental challenge for systematic discovery, as exhaustive experimental screening is infeasible. To address this, we developed an integrated AI-driven framework that combines peptide generation, activity prediction, and toxicity filtering (**Figure 1A**). Our model was trained on a comprehensive, curated dataset of 32,095 peptides (5 to 30 amino acid residues in length), compiled from major AMP databases including APD ^14–17^, dbAMP ^18–20^, DRAMP ^21–24^, dbaasp ^25–27^, CAMP ^28–30^, LAMP ^31,32^, ParaPep ^33^, phytAMP ^34^, AVPdb ^35^, CancerPPD ^36^, AntiTbPdb ^37^, Hemolytik ^38^, and milkAMP ^39^, as well as the training datasets of 24 AMP classifiers, including iAMPCN ^40^, ADAM ^41^, iAMP-2L ^42^, AMPfun ^43^, iAMP-CA2L ^44^, MLACP ^45^, AntiCP 2.0 ^46^, ACPred ^47^, ACPred-FL ^48^, ACPred-Fuse ^49^, ACP-DL ^50^, iACP-DRLF ^51^, mACPpred ^52^, AntiFP ^53^, AVPpred ^54^, AVPIden ^55^, AntiTbPred ^56^, AtbPpred ^57^, BIOFIN ^58^, BIPEP ^59^, dPABBs ^60^, HemoPI ^61^, HLPpred-Fuse ^62^, iAntiTB ^63^, AnOxPePred ^64^, and HAPPENN ^65^.

**Figure 1.**
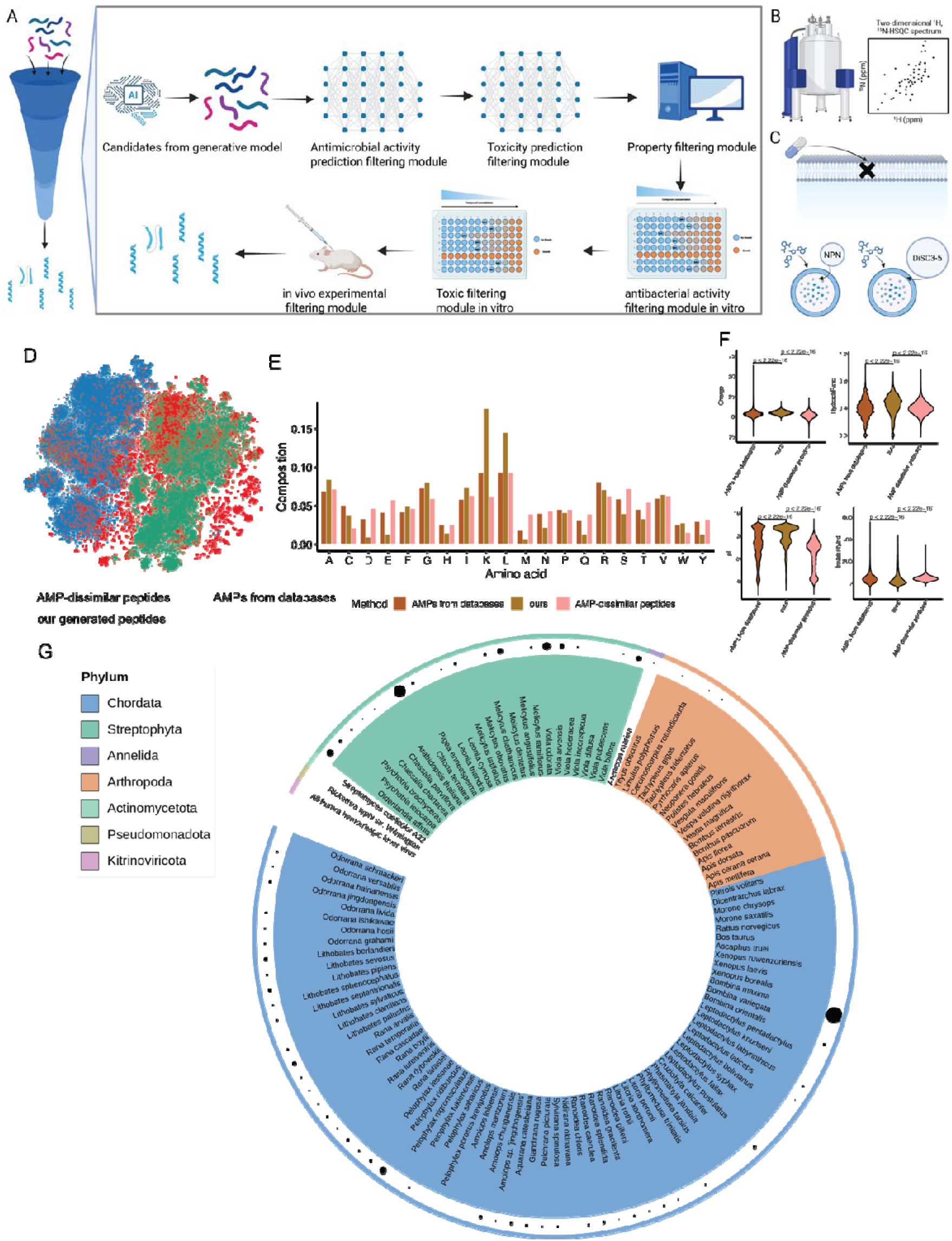
Pipeline for potential AMP development and sequence-related features of generated peptide candidates. **(A)** Schematic overview of the pipeline for mining AMPs from sequences generated by diffusion models, integrating activity prediction, toxicity filtering, property analysis, and experimental validation. **(B)** Secondary structure analysis of representative peptide candidates. **(C)** Proposed antibacterial mechanisms of action, including membrane permeabilization assays. **(D)** t-SNE visualization of AMPs from databases, AMP-dissimilar peptides, and generated peptide candidates. **(E)** Amino acid composition of peptides. Both AMPs from databases and our generated peptides showed enrichment of lysine (K) and leucine (L), whereas acidic residues (D, E) were relatively underrepresented compared with AMP-dissimilar peptides. **(F)** Distribution of physicochemical properties. Violin plots show charge, hydrophobic ratio (the proportion of A, C, F, I, L, M, and V residues), isoelectric point (pI), and instability index across the three peptide groups. **(G)** Taxonomic distribution of proteins or peptides most similar to the generated peptide candidates, covering diverse species from multiple phyla. The size of black dots represents the number of similar sequences, with the strongest similarities observed in *Bombina* (Chordata), and followed by notable matches in *Clitoria* (Streptophyta). Panels **A-C** were created using BioRender.com.

Using Termini, a diffusion-based generative model (“**Diffusion-based generative model for peptide design**” in **Methods**), we generated 1,000 candidate peptides for each length, producing an initial library of 26,000 sequences. This library was subsequently refined through a multi-tier computational screening process. Specifically, we applied a series of predictive models to estimate broad-spectrum antimicrobial activity and to predict minimum inhibitory concentration (MIC) values, and we used a dedicated toxicity classifier to exclude candidates with elevated toxicity risk. This stepwise filtering narrowed the candidate pool to a focused set of leads exhibiting high predicted activity and low risk of toxicity. Finally, the top-ranking candidates were synthesized for experimental validation. Using *in vitro* assays, we assessed antimicrobial activity and toxicity, advancing only those peptides with strong activity and minimal toxicity for *in vivo* evaluation. Mechanistic characterization of representative hits further revealed defined α-helical/β-sheet secondary structures and confirmed membrane-disrupting activity as a primary mode of action (**Figure 1B–C**), linking structure to biological function.

### Generated peptides resemble natural AMPs in sequence, properties, and evolutionary signatures

AMP activity is shaped largely by amino acid composition and physicochemical properties that influence stability and membrane interactions. To assess whether our AI-generated peptides recapitulate the defining characteristics of natural AMPs, we compared generated peptides with AMPs from curated databases and with AMP-dissimilar peptides. We analyzed their sequence distributions using embedding-based visualization (**Figure 1D**), amino acid composition (**Figure 1E**), key physicochemical properties (**Figures 1F and S1**), and sequence similarity to natural protein fragments across species (**Figure 1G**).

A t-SNE visualization based on embeddings from a pretrained ESM protein language model revealed that the generated peptides form a tight cluster with known AMPs, whereas the AMP-dissimilar peptides were distinctly separated (**Figure 1D**). This indicates that our generative model successfully sampled from sequence regions enriched for antimicrobial features rather than generating nonspecific peptides. Further analysis of amino acid composition revealed characteristic AMP-like signatures. Both database and generated AMPs showed enrichment of lysine (K) and leucine (L)—residues critical for conferring with cationic and amphipathic properties—and a marked underrepresentation of acidic residues (aspartic acid, D; glutamic acid, E) compared to the control group, i.e., AMP-dissimilar peptides (**Figure 1E**). An increased frequency of arginine (R) was also observed in the generated peptides, consistent with its crucial role in facilitating electrostatic interactions with negatively charged bacterial membranes.

To evaluate the physicochemical properties of the generated peptides, we compared eight key global properties—net charge, hydrophobicity, isoelectric point (pI), instability index (calculated from dipeptide composition using empirically derived instability weight values), aromaticity (defined as the fraction of aromatic residues, phenylalanine (F), tryptophan (W), and tyrosine (Y)), aliphatic index, Boman index, and hydrophobic ratio [defined as the proportion of alanine (A), cysteine (C), F, isoleucine (I), L, methionine (M), and valine (V) over other residues of the sequence]—against those of database AMPs and AMP-dissimilar peptides (**Figures 1F and S1**). The analysis revealed that the generated peptides exhibited higher net charge and pI values than AMP-dissimilar peptides, with distributions that largely overlapped with those of natural AMPs from databases. This profile reflects an enrichment of cationic residues, a well-established determinant of antimicrobial activity ^66^. In contrast, the AMP-dissimilar peptides exhibited markedly lower charge and reduced isoelectric point (pI), consistent with their lack of antimicrobial properties. Furthermore, the hydrophobicity profiles of the generated peptides closely aligned with those of AMPs from databases, whereas AMP-dissimilar peptides showed a broader distribution with more extreme values. The generated peptides also encoded more stable sequences, as indicated by their lower instability index values, which mirrored the stability of natural AMPs. Additional descriptors further corroborated this AMP-like physicochemical signature. The aromaticity and Boman index values of the generated peptides showed strong overlap with AMPs, whereas the distributions of AMP-dissimilar peptides were either broader or shifted. Although the generated peptides exhibited a slightly higher aliphatic index than database-derived AMPs, indicating a greater contribution of aliphatic side chains, their hydrophobic ratio was more consistent with that of natural AMPs than with non-active, AMP-dissimilar peptides. Collectively, these analyses demonstrate that our generative framework effectively guided peptide design into physicochemical space characteristic of known AMPs, while avoiding the property ranges associated with AMP-dissimilar peptides.

To assess the evolutionary relevance of the generated peptides, we compared their sequences against natural protein fragments across diverse species. The sequence similarity analysis revealed extensive matches to bioactive peptide segments, with the highest number of hits observed in *Bombina maxima* (Chordata), a species renowned for producing skin-derived host-defense peptides. Significant matches were identified in *Clitoria ternatea* (Streptophyta), a plant known for its abundance of defense-related proteins, as well as in several other taxa (**Figure 1G**). The presence of these similarities across both animal and plant lineages suggests that the generated peptides converge upon motifs that have been independently selected in nature to combat microbial threats. Together, these analyses indicate that the framework not only reproduces broad AMP-like features but also generates candidates consistent with motifs observed in natural defense systems across phylogenetically distant clades.

### Predictive and regression models for antibiotic activity

To efficiently screen the vast numbers of generated peptide sequences, we developed a two-tiered predictive modeling approach. First, we constructed binary classification models to predict antimicrobial activity (active vs. inactive) and regression models to estimate potency via log(MIC+1) values (“**Classification models for predicting peptide antibacterial activity**” and “**Regression models for predicting peptide antibacterial activity**” in **Methods; Figure 2A**). Both models were implemented in the Chemprop framework, which employs directed message-passing graph neural networks (GNNs) trained on simplified molecular-input line-entry system (SMILES) representations of the peptides. For the binary classification task, peptides with a minimum inhibitory concentration (MIC) ≤32 µg mL^−1^ against a given bacterial species were labeled as active, while those with higher MICs were considered inactive. This enabled the training of species-specific classifiers to identify candidates with a high likelihood of efficacy. The regression models were trained on the same curated dataset of peptide–MIC pairs to provide a complementary, quantitative estimate of antimicrobial potency.

**Figure 2.**
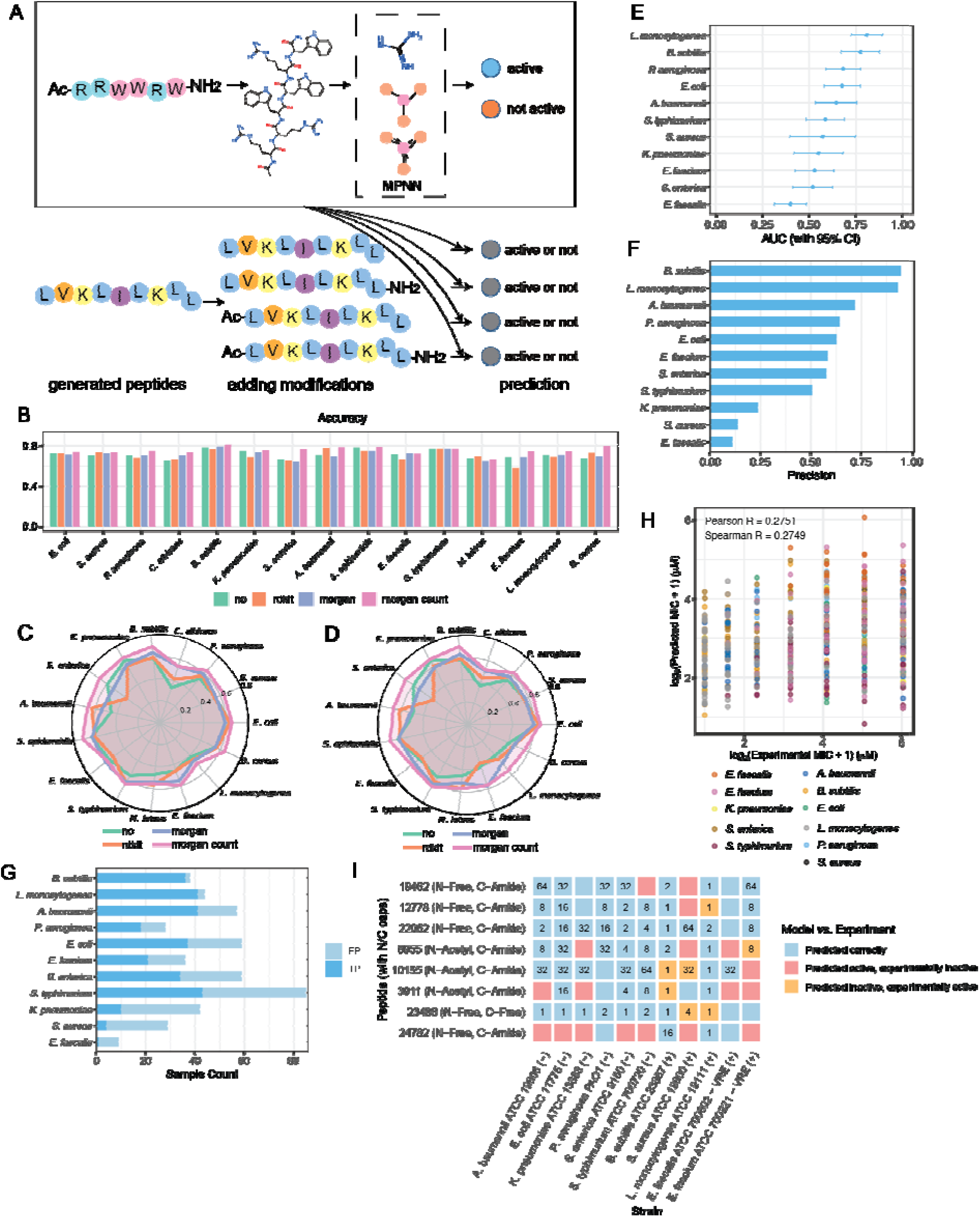
Deep learning framework and evaluation of antibiotic activity predictions using curated datasets and experimental assays of synthesized antimicrobial peptides. **(A)** Schematic of the workflow. Generated peptide sequences were screened with ensembles of models implemented in the Chemprop framework, which is based on graph neural networks (GNNs) and trained on SMILES representations of compounds. The training data were derived from curated experimental datasets, which included peptides with N-terminal acetylation and C-terminal amidation, thereby enabling the models to capture terminal modifications and structural diversity. When applying the models as a filter, these N- and C-terminal modifications were also added to the original sequences to account for capped peptide variants. High-scoring candidates were then prioritized for downstream experimental testing. **(B)** Prediction accuracies of models for predicting peptide activity, evaluated on a randomly selected 20% test set from the curated dataset. **(C–D)** Radar plots showing regression performance, including **(C)** Pearson and **(D)** Spearman correlations, evaluated on an independently selected 20% test set from the same curated dataset. **(E)** Experimental performance of synthesized peptide sequences, measured as AUC with 95% confidence intervals, where one representative strain was used to test predictions at the species level. **(F)** Precision of model predictions evaluated against experimental data from synthesized peptide sequences, using a predictive probability decision threshold of 0.8. **(G)** Experimental validation of synthesized peptide sequences, showing model-predicted active annotations compared with experimentally determined true positive and true negative outcomes. **(H)** Performance of the regression model, comparing predicted versus experimental values for synthesized peptides. **(I)** Experimental performance of predicted broad-spectrum peptides active against multiple species, where one representative strain was used to test predictions at the species level. The color legend indicates the concordance between model predictions and experimental outcomes: predicted correctly (blue), predicted active but experimentally inactive (red), and predicted inactive but experimentally active (yellow). For all experimental evaluations **(E–I)**, species-level prediction performance was tested using one representative strain per species, based on laboratory assays of synthesized peptide sequences.

The predictive models were first evaluated on curated strain-level activity datasets via 10-fold cross-validation and a held-out 20% test set. Ensembles of Chemprop classifiers augmented with Morgan fingerprint features consistently yielded superior performance across all target species (**Figures 2B, S2-3; Tables S1, S2**). In the 10-fold cross-validation, the models achieved AUC values exceeding 0.80 for 9 out of the 15 species (**Figure S2; Table S1**). Performance remained strong for the rest of the six species, with *L. monocytogenes*, *E. faecalis*, *A. baumannii*, *K. pneumoniae*, and *C. albicans* attaining AUCs of 0.75–0.80, while *E. faecium* showed the lowest but still remained in the 0.70–0.75 range.

The high performance was confirmed on the held-out 20% test set. All 15 species achieved AUCs ≥0.75 using the Chemprop classifiers + Morgan count features (**Figure S3**), which consistently outperformed all other alternative feature representations, as measured by both AUC and accuracy (**Figures 2B, S3; Table S2**). High discriminative power (AUCs >0.80) was observed for eight species: *B. subtilis* (0.87), *S. epidermidis* (0.85), *B. cereus* (0.85), *S. enterica* (0.84), *A. baumannii* (0.84), *S. aureus* (0.81), *K. pneumoniae* (0.81), and *P. aeruginosa* (0.80). The remaining seven species showed intermediate AUCs of 0.75–0.80, with *M. luteus* (0.75) representing the lower bound at 0.75, a value still indicative of meaningful predictive ability.

Accuracy metrics generally mirrored the AUC trends, albeit with greater interspecies variability (**Figure 2B**). Two species reached accuracies ≥0.80, including *B. subtilis* (0.81) and *B. cereus* (0.80), while seven species achieved accuracies between 0.75-0.80, including *S. epidermidis* (0.79), *A. baumannii* (0.79), *S. typhimurium* (0.78), *S. enterica* (0.77), *K. pneumoniae* (0.76), *L. monocytogenes* (0.75), and *P. aeruginosa* (0.75). Another five species, such as *E. faecium* (0.75), *E. coli* (0.74), *C. albicans* (0.74), *S. aureus* (0.74), and *E. faecalis* (0.72), achieved accuracies between 0.70 and 0.75. The accuracy for *M. luteus* was the lowest at 0.66; however, its corresponding AUC of 0.75 indicates that the model still retained effective ranking capability. This suggests that applying a more stringent classification threshold (e.g., ≥0.8) could ensure reliable filtering of active candidates for this species. The consistent performance gain from incorporating Morgan fingerprints underscores the added value of explicit molecular feature descriptors for peptide activity prediction. In addition to AUC and accuracy, high precision values across species (**Table S2**) further support the models’ utility for reliably prioritizing candidates in the AMP discovery pipeline. Additional performance evaluation metrics, including recall and F1 scores, are provided in **Table S2**.

The regression models for predicting MIC values were evaluated on curated species-level datasets (with MIC values averaged across strains within each species) using 10-fold cross-validation and a 20% held-out test set. The model incorporating Morgan count fingerprints consistently achieved the lowest average mean squared error (MSE) across species, indicating superior predictive performance (**Figure S4; Table S3**). On the 20% held-out test set, the regression models achieved Pearson correlation coefficients of 0.50–0.72 and Spearman correlations of 0.51–0.71 between predicted and observed log(MIC+1) values (**Figure 2C-D, Table S4-S5**). The highest correlations were observed for S*. epidermidis* (Pearson 0.72; Spearman 0.71) and *B. subtilis* (Pearson 0.70; Spearman 0.70). Strong correlations (0.62–0.70) were also achieved for several key pathogens, including *S. enterica* (Pearson 0.68; Spearman 0.68), *A. baumannii* (Pearson 0.67; Spearman 0.68), *K. pneumoniae* (Pearson 0.68; Spearman 0.68), E. coli (Pearson 0.66; Spearman 0.65), *S. aureus* (Pearson 0.64; Spearman 0.65), *S. typhimurium* (Pearson 0.64; Spearman 0.64), and *P. aeruginosa* (Pearson 0.64; Spearman 0.63). Moderate correlations (0.5–0.6) were obtained for the remaining species, namely *E. faecium*, *E. faecalis*, *M. luteus*, *B. cereus*, *L. monocytogenes*, and *C. albicans*. These correlation levels demonstrate that the regression models captured meaningful quantitative trends in antimicrobial potency. As a result, they can be used as a valuable complementary tool to the classification models, providing an additional filter to rank and prioritize candidate peptides based on their predicted efficacy before experimental validation.

### Antimicrobial activity across a multi-species panel

We evaluated a set of synthesized peptide candidates comprising 120 peptide entries, representing 60 unique backbone sequences with distinct N-terminus (Free vs Acetyl) and C-terminus (Free vs Amide) states, against a panel spanning Gram-negative (*A. baumannii* ATCC 19606, *E. coli* ATCC 11775, *K. pneumoniae* ATCC 13883, *P. aeruginosa* PAO1, *S. enterica* ATCC 9150, and *S. enterica* Typhimurium ATCC 700720) and Gram-positive species (*B. subtilis* ATCC 23857, *S. aureus* ATCC 12600, *L. monocytogenes* ATCC 19111, *E. faecalis* ATCC 700802, and *E. faecium* ATCC 700221) (**Figure S5**). MIC values in this dataset span 1–64 μmol L^−1^. Across the peptides with at least one reported MIC value (111/120 entries), 89/120 showed activity with MIC ≤16 μmol L^−1^ against at least one organism, and 100/120 reached MIC ≤32 μmol L^−1^ against at least one organism.

Assay coverage varied by organism in this dataset (**Figure S5**). Among the most extensively tested species, a high fraction of peptide entries reached MIC ≤16 μmol L^−1^ against *B. subtilis* (79/99 tested) and *L. monocytogenes* (74/88 tested), and against Gram-negative *S. enterica* (52/65 tested) and *A. baumannii* (51/76 tested). In contrast, *S. aureus* ATCC 12600 was measured for a smaller subset (12 peptides), and in that subset only 1/12 reached MIC ≤16 μmol L^−1^. These trends are consistent with broad activity for many candidates in the panel while also highlighting organism-dependent sensitivity and incomplete testing coverage for some strain–peptide combinations (**Figure S5**).

### Broad-spectrum candidates and potency highlights

A substantial subset of peptide entries exhibited broad-spectrum activity in this dataset, defined here as achieving MIC ≤16 μmol L^−1^ against at least one Gram-negative and at least one Gram-positive organism. By this criterion, 63/120 peptide entries meet broad-spectrum activity (**Figure S5**). Several candidates displayed particularly strong and consistent activity across the organisms assayed. For example, peptide 23488 (sequence RRGRFRRAIRRVGRIVVRLVGG, Free N-terminus / Free C-terminus) showed uniformly low MICs across its tested panel (minimum 1 μmol L^−1^; maximum 4 μmol L^−1^ across 9 organisms reported), indicating unusually consistent potency within the measured set (**Figure S5**). Other broadly active entries with wide organism coverage included peptide 9644 (sequence KFKAKLLKKFFKQFKKFL, Free/Free; 11 organisms reported; median MIC 4 μmol L^−1^), and peptide 4194 (sequence WQWRVRLRINKVLPGR, Free/Amide; 11 organisms reported; median MIC 8 μmol L^−1^) (**Figure S5**). Collectively, these results identify multiple candidates with broad activity patterns and highlight specific peptide entries that combine breadth with low MIC values across the assayed organisms (**Figure S5**).

### Predictive performance on the in-house experimental dataset

We next benchmarked the performance of our classification models, which incorporated Morgan count fingerprints, against a validation set of experimentally synthesized peptides. For this evaluation, a single representative strain was selected for each of 11 clinically relevant species: *E. coli*, *S. aureus*, *P. aeruginosa*, *B. subtilis*, *K. pneumoniae*, *S. enterica*, *A. baumannii*, *E. faecalis*, *S. typhimurium*, *E. faecium*, and *L. monocytogenes*. To make a comparative analysis, regression model outputs predicting log(MIC+1) > 5 were assigned a predictive probability of 0 prior to evaluation. At the species level, models incorporating Morgan count fingerprints achieved AUC values exceeding 0.6 for six species, with two species demonstrating particularly strong predictive performance: *L. monocytogenes* (0.81, 95% CI: 0.73–0.89) and *B. subtilis* (0.77, 95% CI: 0.67–0.88) (**Figure 2E; Table S6**). In contrast, the lowest discriminative ability was observed for *E. faecalis* (0.40, 95% CI: 0.31–0.49) (**Figure 2E; Table S6**).

Critically, even for species with low overall AUCs such as *E. faecalis*, *S. enterica*, and *E. faecium*, the models successfully recovered true positives when a stringent probability cutoff (>0.8) was applied. For example, this filtering yielded one active hit for *E. faecalis* (precision = 0.11), 34 active hits for *S. enterica* (precision = 0.58), and 21 active hits for *E. faecium* (precision = 0.58) (**Figure 2F, 2G; Table S7**). These results underscore the practical utility of our models for species-level screening. By effectively enriching large candidate pools for true positives, even recovering valuable hits in challenging cases, the developed framework can significantly reduce experimental burden and accelerate the discovery of promising therapeutic candidates against specific pathogens at the species level.

### Regression performance and broad-spectrum candidate identification on the in-house experimental dataset

To evaluate regression performance on experimental data, we also tested the MIC assays of the synthesized peptide sequences against 11 bacterial strains, each representing a distinct species. The overall correlation between predicted versus experimentally measured MIC values was moderate, with Pearson and Spearman correlation coefficients of 0.28 and 0.27, respectively (**Figure 2H**). We further assessed the framework’s utility as a predictive filter by applying a stringent decision threshold of 0.8 to laboratory outcomes. Across the 11 species, the model effectively enriched for active candidates, yielding 386 true-positive hits from 1,320 prediction–experiment pairs. Under the same experimental activity cutoff (MIC ≤ 64 μmol L^−1^), the strain-specific APEX model identified 146 true positives from 536 prediction–experiment pairs based on its 69 synthesized peptides. Together, these results support the utility of our framework as a scalable high-throughput screening tool for prioritizing active antimicrobial candidates (**Figure 2G; Table S7**).

Performance varied notably by species. For *B. subtilis* (36 TP vs. 2 FP; precision = 0.95) and *L. monocytogenes* (41 TP vs. 3 FP; precision = 0.93), the model demonstrated exceptionally high precision, strongly enriching for true actives. Substantial enrichment was also observed for *E. coli* (37 TP vs. 22 FP) and *P. aeruginosa* (18 TP vs. 10 FP). In contrast, predictive filtering was less effective for *S. aureus* (4 TP vs. 25 FP; precision = 0.14) and *E. faecalis* (1 TP vs. 8 FP; precision = 0.11). Nevertheless, even for these challenging species, the precision significantly exceeded typical hit rate of random peptide screening (1-2%), substantially reducing the experimental burden.

We also evaluated the model’s ability to identify broad-spectrum candidate peptides. Eight peptides predicted to be active against multiple bacterial species were selected for validation (**Figure 2I**). Experimental assays confirmed that most of these peptides indeed exhibited broad-spectrum activity. For instance, the peptides 19462 (N-Free, C-Amide) and 12778 (N-Free, C-Amide) were predicted to be active across nearly all species and demonstrated experimental growth inhibition against *A. baumannii*, *E. coli*, *K. pneumoniae*, *P. aeruginosa*, *S. enterica*, *B. subtilis*, and *E. faecalis*. Similarly, the peptides 10155 (N−Acetyl, C−Amide) and 22062 (N-Free, C-Amide) showed extensive cross-species activity across both Gram-positive and Gram-negative strains. Only one peptide, 24782 (N−Acetyl, C−Amide), was consistently predicted as active but failed to show experimental efficacy. Overall, the concordance between predicted and experimentally validated activities was high, with agreement in 63 out of 88 prediction–experiment pairs (71.6%) across species. Notably, seven of the eight synthesized peptides exhibited activity against three or more bacterial species, while only one peptide was restricted to activity against two species. These results underscore the framework’s utility in discovering broad-spectrum antimicrobials.

### Impact of terminal modifications on measured antibacterial activity

Because multiple backbone sequences were tested in more than one terminal state (Free vs Acetyl N-terminus; Free vs Amide C-terminus), we examined how terminal modifications relate to the measured MIC patterns within this dataset. For several backbones where both Free/Free and Free/Amide variants were measured on overlapping organisms, C-terminal amidation was frequently associated with lower MICs across the shared test set (**Figure 3A**). Notably, this trend was observed for backbones such as FKKAIKKWSKSWAPWWK (Free/Amide: peptide 7374; Acetyl/Amide: peptide 7375; **Figure S5**) and RRRIKWRRGAIRPAVIRLVKSV (Free/Amide: peptide 22062; Acetyl/Amide: peptide 22063; **Figure 3A**), each of which displayed broad-spectrum activity with multiple MICs ≤16 μmol L^−1^ across the organisms tested. At the same time, the magnitude and direction of terminal effects were not uniform across all backbones represented here, underscoring that terminal capping can measurably shift activity but does so in a sequence-dependent manner (**Figure 3A**).

**Figure 3.**
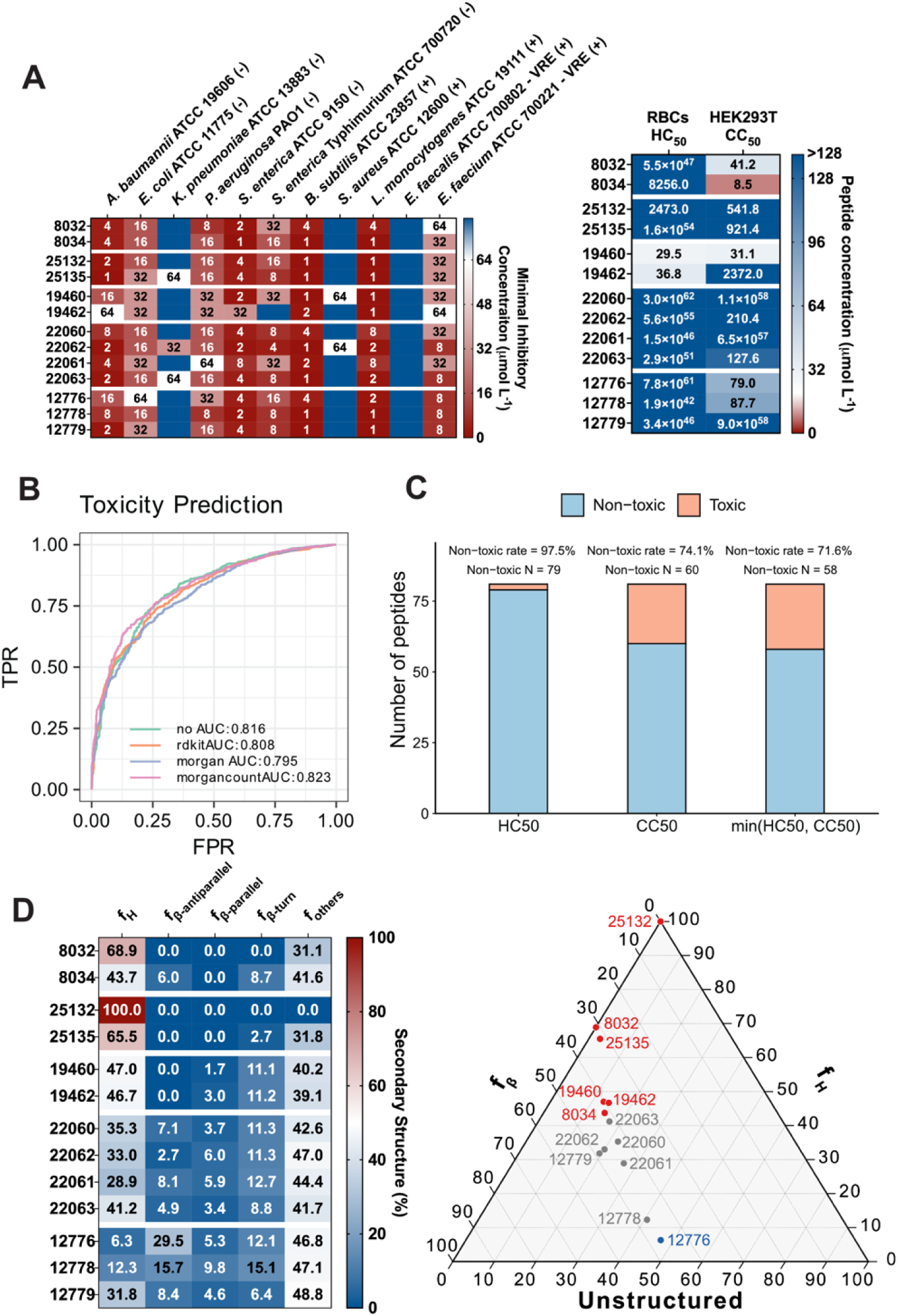
Antimicrobial, cytotoxic and hemolytic activity, performance of toxicity prediction models, consistency of modification effects, and secondary structure of selected peptides. **(A)** Heat map of minimum inhibitory concentrations (MICs, μmol[L^−1^) against 11 clinically relevant Gram-negative (–) and Gram-positive (+) pathogens, including antibiotic-resistant strains. Bacteria (10^5^ CFU) were incubated with serial peptide dilutions (0–64 μmol□L^−1^) at 37□°C, and growth was quantified by OD_600_ after 24 h. MIC values represent the mode of replicate measurements for each condition. Hemolytic (HC_50_) and cytotoxic (CC_50_) concentrations, defined as the peptide concentration causing 50% lysis of red blood cells or reduction in HEK293T cell viability, respectively. Values were obtained by nonlinear regression of dose-response curves. Data represent three independent experiments. **(B)** Prediction accuracies of models for predicting peptide toxicity, evaluated on a randomly selected 20% test set from the curated dataset. **(C)** Experimental validation of the toxicity predicted score as a counter-screening filter. Of the 81 peptides selected using a toxicity score threshold of 0.2, 97.5% were non-hemolytic based on HC_50_, 74.1% were non-cytotoxic based on CC_50_, and 71.6% were classified as non-toxic using the combined endpoint min(HC_50_, CC_50_). Peptides with HC_50_ or CC_50_ values > 64 μmol L^−1^ (estimated by non-linear regression) were considered non-hemolytic or non-cytotoxic, respectively. **(D)** Heat map shows the percentage of secondary structure for each peptide calculated using the BeStSel algorithm. Ternary plot showing the secondary structure fraction for each of the selected peptides.

### Directional consistency of predicted and experimental modification effects

We next evaluated whether the model’s predicted changes in activity following terminal modification were directionally consistent with the corresponding experimental MIC changes across bacterial strains. This directional agreement was visualized in a heatmap (**Figure S6**). Across all comparisons, we observed 60 matches, 28 mismatches, and 37 neutral outcomes. Neutral outcomes occurred when the experimental MIC values for the modified and unmodified peptides were identical. This is an inherent limitation of the broth microdilution method, which used two-fold serial dilutions; when both variants fall within the same dilution step, the true direction of change cannot be determined. Accordingly, these neutral cases were excluded from the performance evaluation. Direction matches were frequently observed across species, with 68% (60/88) matches and 32% (28/88) mismatches. This indicates that the predictive models captured the relative impact of N- or C-terminal modifications on antibacterial potency. This capability was further illustrated by analyzing two specific peptides (22060 and 24780). For peptide 22060, the model achieved a match rate of 86.4% (19 matches vs. 3 mismatches) across all modification variants. In contrast, for peptide 24780, all experimental MIC values for its variants fell within the same dilution step (yellow cells in the heatmap), precluding a directional assessment. Collectively, these results demonstrate that our predictive model can accurately capture the directionality of activity changes induced by terminal modifications. This provides a powerful, biologically meaningful signal for prioritizing the most promising peptide variants during experimental screening.

### Predictive models for peptide cytotoxicity

To complement our antibiotic activity predictions and enable the selection of candidates with low toxicity, we constructed cytotoxicity prediction models using the same Chemprop framework (**Figure 2A**; “**Toxicity prediction models**” in **Methods**). Peptides were represented in SMILES format, and the models were trained to classify them as cytotoxic or non-cytotoxic based on curated datasets.

We systematically evaluated the cytotoxicity prediction models using 10-fold cross-validation and a held-out 20% test set. Among the various feature representations, Chemprop classifiers augmented with Morgan count fingerprints achieved the best overall performance, attaining the highest average AUC of 0.814 in cross-validation and 0.823 on test set (**Figure 3B, S7; Table S8**). Consequently, this model was selected as the final classifier for cytotoxicity filtering of peptides.

We next evaluated the toxicity score as an experimental counter-screening filter using our 120 synthesized peptides. Based on experimental measurements, peptides with half maximal hemolytic concentration (HC_50_) or half maximal cytotoxic concentration (CC_50_) values greater than 64 μmol L□¹ (estimated by non-linear regression) were considered non-toxic, while the remaining peptides were classified as toxic. Rather than treating cytotoxicity prediction as a standalone classification task, we primarily assessed whether the toxicity score could enrich non-toxic candidates among peptides selected for experimental testing.

We next applied the predictive toxicity score as a counter-screening filter to triage peptides for experimental validation. Using a threshold of 0.2, 81 peptides were selected from the designed candidates and subjected to cytotoxicity assays. Experimental toxicity was then evaluated based on measurements of the HC_50_ and the CC_50_. 79 of the 81 selected peptides (97.5%) were experimentally confirmed as non-hemolytic as assessed by HC50 (**Figure 3C; Table S9**). 60 of the 81 selected peptides (74.1%) were experimentally non-cytotoxic as assessed by CC50 (**Figure 3C; Table S9**). Finally, under the combined endpoint defined as min(HC50, CC50), 58 of the 81 selected peptides (71.6%) were experimentally classified as non-toxic (**Figure 3C; Table S9**). Collectively, these results demonstrate that the toxicity score is particularly effective at filtering hemolytic peptides, while also providing measurable enrichment against general cytotoxicity. These findings support the intended use of the toxicity score as an experimental triaging filter, rather than as a definitive cytotoxicity classifier.

### Experimental cytotoxicity and hemolysis profiling of synthesized candidates

We next assessed mammalian cell compatibility and red blood cell lysis for the same synthesized peptide set used for antimicrobial testing by measuring cytotoxic (CC_50_; HEK293T cells) and hemolytic (HC_50_; red blood cells) concentrations needed to kill 50% of the cells (**Figure S8**). Across the 120 peptide entries, CC_50_ values ranged from 6.5 μmol L^−1^ to values exceeding the upper assay limit (128 μmol L^−1^), indicating that the panel spans from cytotoxic peptides to peptides with minimal detectable cytotoxicity under the tested conditions (**Figure S8**). HC_50_ values similarly ranged from 0.15 μmol L^−1^ to values at or beyond the assay limit (128 μmol L^−1^), reflecting wide variability in hemolytic propensity across the library. Using the binary toxicity definition applied for model validation (CC_50_ > 64 μmol L^−1^ as non-toxic), 87/120 peptides were labeled non-toxic and 33/120 peptides were labeled toxic (**Figure S8**).

Overall, cytotoxicity and hemolysis were related but not redundant (**Figure S8**). While many peptides with high CC_50_ also showed little measurable hemolysis within the assay window, several sequences deviated from this coupling, underscoring that mammalian cytotoxicity and red blood cell lysis capture overlapping but distinct liabilities. For instance, the backbone PWKLWKKVIKLVKKWLRL exhibited high hemolysis in both tested terminal forms (HC_50_ of 0.1797 μmol L^−1^ for Free/Free, peptide 10152, and HC_50_ of 0.1524 μmol L^−1^ for Acetyl/Amide, peptide 10155), yet the corresponding CC_50_ values differed markedly between variants (CC_50_ of 9.11 μmol L^−1^ for Free/Free versus an assay-limit CC_50_ for Acetyl/Amide), illustrating that terminal chemistry can decouple cytotoxic and hemolytic behavior for the same backbone sequence.

Because multiple backbones were measured in more than one terminal state, we examined whether N-terminal acetylation and/or C-terminal amidation systematically shifted cytotoxicity and hemolysis for identical sequences. These effects were clearly sequence-dependent rather than uniform across the library. A particularly informative example is the four-variant backbone RRFIKRGKTAIRRRPRAIIIRII, which was tested as Free/Free (peptide 24780), Free/Amide (24782), Acetyl/Free (24781), and Acetyl/Amide (24783). In this case, N-terminal acetylation without C-terminal amidation improved the safety profile relative to the unmodified peptide: Acetyl/Free increased CC_50_ to 120.50 μmol L^−1^ compared to 28.78 μmol L^−1^ for Free/Free, while also maintaining high HC_50_ values (HC_50_ of 6167 vs 21935, respectively). In contrast, for the same backbone, both amidated variants (Free/Amide and Acetyl/Amide) showed lower CC_50_ values (35.60 and 24.44 μmol L^−1^, respectively), indicating that amidation did not universally reduce cytotoxicity for this sequence.

Other backbones demonstrated the opposite behavior, where terminal changes increased toxicity. For example, for KKWKKFFKAAKKFAKKIG, C-terminal amidation substantially decreased both CC_50_ and HC_50_ (Free/Free: CC_50_ of 41.19, HC_50_ recorded at assay limit; Free/Amide: CC_50_ of 8.481, HC_50_ of 8256), consistent with a shift toward increased mammalian toxicity in the amidated form for this backbone. Similarly, FTGRWKWRSRFRKRRWYT showed reduced CC_50_ upon N-terminal acetylation (Free/Free CC_50_ of 90.32 vs Acetyl/Free CC_50_ of 22.47), whereas PVVRRFRWWRSFRIRR displayed improved cytotoxicity in acetylated variants (Free/Free CC_50_ of 132.8 vs Acetyl/Free CC_50_ of 361.5 and Acetyl/Amide CC_50_ of 315.1). Together, these paired comparisons across matched backbones indicate that terminal chemistry can strongly influence mammalian compatibility, but the direction and magnitude of the effect depend on the underlying sequence context.

Integrating toxicity readouts (**Figure S8**) with the antimicrobial results (**Figure S5**) highlights that several candidates achieve antibacterial potency while maintaining favorable CC_50_ and HC_50_ profiles, supporting the rationale for using the cytotoxicity classifier as a pre-synthesis filter. At the same time, the observed sequence-dependent terminal effects emphasize why explicitly representing terminal modifications during both generation and screening is important: identical backbones can shift from non-toxic to toxic (or vice versa) with capping changes, and the same modification can improve one backbone while worsening another. These results therefore provide experimental support for a modification-aware design framework and motivate future work to learn modification–sequence interactions more directly within the toxicity prediction task.

### Physicochemical landscape of synthesized peptide candidates

Based on the generative and filtering pipeline (“**Down-selection of generated peptides for synthesis and testing**” in **Methods**), we synthesized 120 peptides representing 60 unique backbone sequences, each produced with and without terminal modifications, to downstream profiling (**Data S1**). Eighty candidates were prioritized by the pipeline, while the remaining 40 were included as a comparative set to assess terminal modification effects. To determine whether these candidates occupy a distinct region of the physicochemical space relative to known AMPs, we compared them against curated AMPs from public databases and with AMP-dissimilar sequences (**Figure S9**). A t-SNE projection revealed that database AMPs formed dense, well-defined clusters, whereas the synthesized peptides localized to peripheral or sparsely populated regions of the embedding, consistent with divergence from previously characterized AMP families (**Figure S9A**). Analyses of key physicochemical features reinforced this separation. The synthesized peptides displayed significantly higher isoelectric points and net positive charges compared than database AMPs (**Figures S9B-S9D**), while the instability index showed no significant differences between groups (**Figure S9C**).

In contrast, synthesized candidates displayed markedly reduced hydrophobic fractions (**Figure S9E**), suggesting that many fall outside the conventional amphipathic balance of hydrophobic and cationicity characteristics of canonical membrane-active AMPs. Consistent with these shifts, amino acid composition analysis showed increased frequencies of K, proline (P), R, and V, alongside a pronounced reduction in M relative to known AMPs (**Figure S9F**). These compositional changes provide a coherent explanation for the observed physicochemical trends: elevated pI and charge reflect enrichment in K/R, whereas reduced hydrophobic ratios are consistent with depleted M content despite a modest increase in V. Together, these results indicate that our candidates constitute a distinct class of sequences with physicochemical profiles characterized by high cationicity, comparatively low hydrophobicity, and altered residue usage, thereby expanding the accessible sequence–property space beyond that represented in current AMP databases (**Figure S9**). Finally, comparative sequence analysis identified no detectable similarity between the synthesized peptides and natural proteins (“**Comparative sequence analysis of generated peptides**” in **Methods**), supporting the novelty of the generated sequences.

### Secondary-structure propensities of synthesized peptides at a hydrophobic–hydrophilic interface

To experimentally probe the folding behavior of the synthesized peptides under membrane-relevant conditions, we performed circular dichroism (CD) spectroscopy in aqueous buffer containing 10 mmol L^−1^ SDS, which provides a well-established hydrophobic–hydrophilic interface that can promote secondary structure formation in responsive sequences (**Figure S10**). CD spectra were analyzed using the BeStSel algorithm to estimate fractional contributions from α-helical conformations (f_H_), β-structures (f_β_; including antiparallel and parallel components as well as turns), and unstructured conformations.

Across the full peptide set, the CD data revealed pronounced heterogeneity in secondary structure rather than convergence toward a single dominant fold. While a subset of peptides exhibited high α-helical content in SDS (e.g., 13742/13743, 25132, 31315, 23513), many populated mixed conformational states, and a substantial fraction remained largely disordered despite the presence of an interfacial mimic (e.g., 4194/4192, 20716, 6892). Peptides with elevated helical fractions generally corresponded to longer sequences enriched in hydrophobic residues interspersed with basic amino acids and showed low proline content, consistent with an intrinsic capacity to form amphipathic helices at interfaces (e.g., 13742/13743, 23513, 25132, 9644). However, these highly helical peptides represented only a minority of the library and did not define the dominant structural behavior across all candidates.

In contrast, a distinct subset of peptides displayed appreciable β-structure content, with antiparallel β components contributing most prominently (e.g., 44339, 1472, 12776, 10413, 4088, 7372). These β-enriched peptides often exhibited moderate to low helical fractions and were not simply unfolded; instead, they populated structured but non-helical conformational ensembles, frequently accompanied by increased β-turn contributions (e.g., 44339, 7524/7526, 20717, 16092). Such behavior was observed across peptides of varying lengths and compositions, indicating that β-structure formation in this dataset is not restricted to a narrow sequence class. Notably, parallel β contributions were consistently low, suggesting that β-structure, when present, is dominated by antiparallel arrangements rather than extended β-sheet assemblies.

A large fraction of the peptide library retained high unstructured fractions in SDS, indicating limited stabilization of regular secondary structure under these conditions (e.g., 4194/4192, 20716, 8952, 7704, 18946). These predominantly disordered peptides were typically characterized by high net positive charge and relatively low hydrophobic residue content, features that favor interfacial association without enforcing stable folding. The prevalence of partially or largely disordered conformations underscores that many candidates interact with membrane-like environments through flexible, adaptive binding modes rather than through pre-organized secondary structures.

Because multiple backbone sequences were examined in more than one terminal state, the CD data also enabled direct evaluation of the influence of N-terminal acetylation and C-terminal amidation on secondary structure. Across the dataset, terminal modifications did not exert a uniform or directional effect on folding. For some backbones, C-terminal amidation was associated with increased α-helical content, consistent with reduced terminal fraying and enhanced helix stabilization (e.g., 19826 → 19824, and 1472 → 1474). For other sequences, the same modification produced little change or even reduced helicity (e.g., 8032 → 8034 – **Figure 3D**, and 18944 → 18946), while in several cases N-terminal acetylation modulated the balance between helical and disordered fractions without inducing substantial β-structure (e.g., 23512 → 23513 – **Figure 3D**, and 10412 → 10413). These observations indicate that terminal chemistry influences folding in a strongly sequence-dependent manner rather than acting as a global switch toward ordered structure (e.g., minimal change in an already highly helical backbone, 13742 ↔ 13743).

Taken together, the CD-derived secondary structure profiles demonstrate that the synthesized peptides span a broad conformational landscape at hydrophobic–hydrophilic interfaces, encompassing highly helical, β-enriched, mixed, and largely disordered states. The absence of systematic convergence toward α-helicity—even under conditions favorable for helix formation—highlights the structural diversity encoded in the library and emphasizes that antimicrobial function in these candidates does not rely on a single canonical fold. Instead, the data support a model in which secondary structure emerges contextually from the interplay between sequence composition, terminal chemistry, and interfacial environment, reinforcing the design principle that biological activity can be achieved through multiple structural solutions.

### Bacterial membrane perturbation revealed by permeabilization and depolarization assays

To resolve how membrane-active peptides differentially affect the bacterial envelope, we quantified both outer membrane permeabilization and cytoplasmic membrane depolarization using NPN and DiSC_3_-5 assays, respectively (**Figure 4A-B**). For each assay, fluorescence kinetics were summarized using maximum relative fluorescence (MaxRel) and area under the curve (AUC), capturing peak intensity and sustained response over time. Within each assay, MaxRel and AUC were tightly coupled, indicating that peptides producing strong peak signals also tended to induce prolonged membrane perturbation. This consistency enabled direct comparison of membrane interaction phenotypes across peptides and between assays.

**Figure 4.**
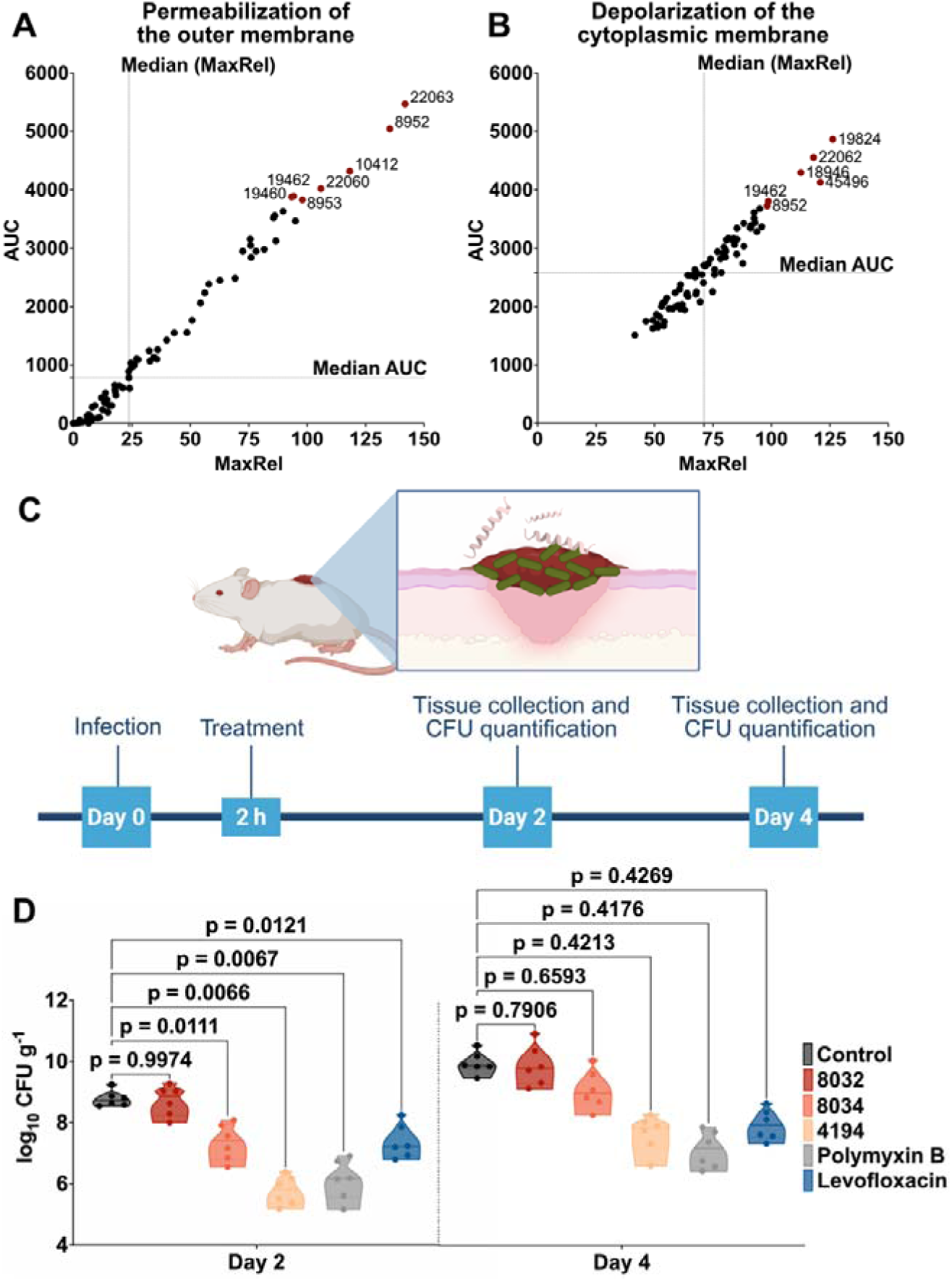
Membrane-targeting mechanism and *in vivo* efficacy of optimized peptides. **(A-B)** Active peptides against *A. baumannii* ATCC 19606 were evaluated for outer and cytoplasmic membrane disruption. **(A)** Outer membrane permeabilization measured by NPN uptake. Each point represents a peptide plotted as maximum relative fluorescence (MaxRel) versus area under the curve (AUC). Dashed lines denote median values used to classify peptides into four response profiles: strong permeabilizers (robust and sustained disruption; upper right), transient permeabilizers (rapid, short-lived response; upper left), slow permeabilizers (gradual fluorescence increase; lower right), and weak permeabilizers (minimal activity; lower left). Representative peptides are indicated. **(B)** Cytoplasmic membrane depolarization assessed using DiSC[-5. Peptides are plotted as MaxRel versus AUC and classified analogously into potent, transient, gradual, and weak depolarizers. Representative potent depolarizers are labeled. **(C)** Schematic of the murine skin abscess model used to evaluate peptides 8032, 8034 and 4194 (n = 6 per group) against *A. baumannii* ATCC 19606. **(D)** A single 10× MIC dose was administered post-infection. Capped peptide 8034 prolonged infection control for two days relative to the wild-type peptide (8032), while capped peptide 4194 maintained suppression for up to four days compared to untreated controls and reference antibiotics (polymyxin B and levofloxacin). Panel **C** was created with BioRender.

Joint analysis of NPN and DiSC_3_-5 responses revealed a continuum of membrane interaction behaviors, rather than a single dominant mechanism. Some peptides induced strong responses in both assays, others showed preferential outer membrane permeabilization or depolarization, and a subset exhibited minimal signal in one or both assays despite measurable antimicrobial activity. These patterns highlight mechanistic diversity within the library and underscore that membrane perturbation is modulated by sequence context and terminal chemistry rather than dictated by a single physicochemical or structural feature.

A subset of peptides produced high MaxRel and AUC values in both assays, consistent with coordinated disruption of the outer membrane followed by loss of cytoplasmic membrane potential. The most prominent example is 22063 (RRRIKWRRGAIRPAVIRLVKSV, Acetyl/Amide), which ranked highest in permeabilization and also among the strongest depolarizers. Importantly, this peptide displays intermediate α-helical content in SDS with a high disordered fraction, indicating that effective depolarization does not require a fully stabilized helical conformation. In line with this, 22063 combines strong Gram-negative activity with favorable hemolysis and cytotoxicity profiles, demonstrating that potent membrane perturbation can be achieved without nonspecific mammalian membrane damage.

A similar coupling was observed for 8952 (RWWRRVRKKFKKNLVFLK, Free/Free), which showed very strong responses while remaining largely disordered in SDS. These examples reinforce the conclusion drawn from the secondary structure experiments that dynamic, partially disordered conformational ensembles can efficiently perturb bacterial membranes and drive depolarization.

In contrast, several peptides showed robust permeabilization responses but comparatively weaker depolarization, indicating that outer membrane permeabilization does not always translate directly into rapid loss of cytoplasmic membrane potential. The backbone FKKAIKKWSKSWAPWWK illustrates this decoupling. The Acetyl/Amide variant (7375) exhibited strong permeabilization alongside improved antimicrobial activity, yet its depolarization response was more moderate than that of peptides such as 22063. Notably, the Free/Amide variant (7374) produced no detectable permeabilization signal but still showed measurable depolarization and retained Gram-negative activity. These observations suggest that, for certain sequences, access to or perturbation of the inner membrane can occur without pronounced outer membrane damage under the assay conditions.

Conversely, a smaller group of peptides displayed clear depolarization despite weak or negligible permeabilization responses, highlighting cases where cytoplasmic membrane perturbation is not preceded by strong outer membrane permeabilization. For example, 41804 (APIFAKILKGGNNAFKNLKKLGLGFKFG, Free/Free) showed minimal permeabilization signal yet induced detectable depolarization and maintained Gram-negative antimicrobial activity. These peptides provide concrete examples that outer membrane permeabilization, as captured by NPN, is not an absolute prerequisite for depolarization or antibacterial efficacy, at least within the temporal and concentration regimes probed here.

Direct comparison of peptides sharing the same backbone but differing in terminal chemistry revealed that N-terminal acetylation and C-terminal amidation can tune both permeabilization and depolarization in a strongly sequence-dependent manner. For RRRIKWRRGAIRPAVIRLVKSV (22060–22063), progressive enhancement of both responses was observed from Free/Free to Acetyl/Amide, occurring alongside only modest changes in secondary structure. This indicates that terminal modifications can modulate membrane interaction kinetics without enforcing a major structural transition.

In contrast, the backbone GLVNIGKKVVGKALGYGAYSASGKLKLVYN (49988–49991) showed generally weak responses across all terminal states, despite maintaining moderate helicity in SDS micelles. Here, acetylation and amidation had only subtle effects on membrane perturbation, reinforcing that helical propensity alone is insufficient to predict either permeabilization or depolarization.

The PWKLWKKVIKLVKKWLRL backbone (10152/10155) highlights a different regime: both terminal variants produced strong permeabilization and depolarization signals and were also strongly hemolytic, indicating that for this sequence, membrane depolarization correlates with reduced selectivity toward bacterial membranes. This contrasts with sequences such as 22063, where strong depolarization is decoupled from mammalian membrane toxicity, underscoring the dominant role of sequence context in determining selectivity.

Taken together, the combined permeabilization and depolarization datasets demonstrate that the peptide library spans multiple membrane interaction strategies, including peptides that (i) strongly permeabilize and depolarize, (ii) primarily permeabilize the outer membrane, (iii) depolarize with limited outer membrane signal, and (iv) exhibit weak responses in both assays yet retain antimicrobial activity. These mechanistic classes cut across secondary structure categories: highly helical, mixed, β-enriched, and largely disordered peptides are represented in each regime, consistent with the secondary structure results showing broad structural diversity in SDS micelles.

Importantly, terminal modifications act as context-dependent modulators of both permeabilization and depolarization rather than as universal switches toward membrane disruption. Together, these results support a mechanistically pluralistic model in which effective antibacterial activity can arise from distinct modes of membrane interaction, rather than a single canonical pathway dominated by stable α-helical pore formation.

### Terminal chemistry repositions peptides within the membrane-disruption process

Terminal modification produced substantial changes in antimicrobial potency, membrane permeabilization and secondary structure across peptide backbones. However, these observables did not vary monotonically with each other. Increased helicity did not consistently correspond to stronger killing, outer-membrane permeabilization did not necessarily coincide with depolarization, and peptides with comparable charge density displayed markedly different MIC values. These inconsistencies indicated that activity could not be explained by any single physicochemical parameter such as folding propensity, hydrophobicity, or electrostatic attraction.

To determine whether these divergent phenotypes reflected changes in binding location or changes in the subsequent membrane-interaction process, we compared experimental measurements with residue-level attribution patterns derived from Chemprop. For each peptide, activity-associated molecular fragments were mapped onto the backbone to recover contributing residues, which were merged into contiguous functional segments (**Table S10**). This allowed direct comparison between the spatial organization of membrane engagement and the experimentally observed permeabilization, depolarization and antimicrobial activity.

For the FKKAIKKWSKSWAPWWK backbone, unmodified and Free/Am variants displayed two spatially separated determinants: an N-proximal cationic/aromatic region (KKW) and a broader mid-to-C segment (KSWAPWW), consistent with diffuse electrostatic adsorption. After acetylation and amidation, attribution collapsed into a single localized segment centered on APW, coinciding with increased outer-membrane permeabilization and improved antimicrobial potency. The emergence of a confined determinant indicates that terminal chemistry enabled formation of a discrete insertion interface rather than merely strengthening association.

In contrast, RWWRRVRKKFKKNLVFLK displayed broadly distributed attributions across the sequence in the Free/Free, Ac/Free, and Ac/Am states, without a dominant hotspot. N-terminal acetylation increased helical propensity while reducing β/turn content, yet the peptide remained largely disordered after combined acetylation and amidation. Outer-membrane permeabilization (NPN) remained in the high-response regime in two states, with only modest rank differences. The combination of distributed determinants, preserved activity, and incomplete structural consolidation supports membrane engagement through multivalent surface contacts rather than a single defined insertion conformation, with terminal chemistry tuning disruption efficiency rather than interaction mode.

For RRRIKWRRGAIRPAVIRLVKSV, attribution consistently localized to an N-proximal cationic anchor at R(1) and the RPAV(12–15) block in the Free/Free, Ac/Free, and Ac/Am states, indicating a conserved binding locus. These observations suggest that terminal chemistry did not redirect the interaction site but enhanced the efficiency of membrane disruption downstream of initial adsorption.

The backbone GLVNIGKKVVGKALGYGAYSASGKLKLVYN (49988–49991) exhibited weak NPN permeabilization and DiSC3-5 depolarization in the Free/Free, Ac/Free, Free/Am, and Ac/Am states, despite maintaining moderate helicity in SDS micelles. Attribution was broadly distributed along the sequence without a dominant hotspot. Together, these observations indicate that, for this sequence context, helical propensity and global folding alone are not sufficient to yield productive membrane disruption, implying that additional features—such as the spatial organization of charge and hydrophobicity or specific residue-level determinants—are required to couple adsorption to downstream perturbation.

Together these cases show that terminal chemistry does not operate along a single activity axis such as charge, helicity or hydrophobicity. Instead, peptides differ in which step of the envelope interaction pathway limits killing: formation of a discrete insertion locus, cooperative surface engagement, or post-binding membrane traversal. Chemprop attribution interpretation helped distinguish changes in interaction topology from changes in interaction strength. The combined data are consistent with a model in which terminal modification shifts peptides along a membrane-interaction pathway by altering the rate-limiting transition rather than relocating the binding site.

### *In vivo* efficacy in a murine skin infection model

To assess *in vivo* efficacy, we evaluated three lead candidates in a murine skin scarification model of *Acinetobacter baumannii* ATCC 19606 infection (**Figure 4C**), a clinically relevant Gram-negative pathogen associated with skin and soft-tissue infections.

Peptides 8032 (KKWKKFFKAAKKFAKKIG, Free/Free), 8034 (KKWKKFFKAAKKFAKKIG, Free/Amide), and 4194 (WQWRVRLRINKVLPGR, Free/Amide) were selected based on complementary *in vitro* profiles. Peptides 8032 and 8034 constitute a matched unmodified/modified pair with identical backbones, enabling direct assessment of C-terminal amidation effects *in vivo*, while peptide 4194 served as a mechanistically and structurally distinct comparator. All three peptides showed activity against *A. baumannii in vitro* and favorable toxicity profiles relative to their antibacterial potency (**Figure 4D**).

At day 2 post-infection, topical treatment with peptides 8032, 8034, and 4194 resulted in a significant reduction in bacterial burden compared with untreated controls (p = 0.0066, 0.0067, and 0.0121, respectively), with efficacy comparable to polymyxin B (p = 0.0111).

By contrast, vehicle treatment showed no effect. At day 4, bacterial loads converged across all groups and no statistically significant differences were detected, consistent with partial resolution of infection under the dosing conditions used.

The comparable *in vivo* efficacy of peptides 8032 and 8034 indicates that C-terminal amidation does not compromise antibacterial performance. Peptide 4194 also produced significant early bacterial clearance. Together, these results demonstrate that peptides prioritized by the pipeline can reduce bacterial burden in a relevant *in vivo* infection model and that terminal modification can tune safety without sacrificing efficacy. The transient nature of the effect highlights opportunities for optimization through dosing and/or formulation, while supporting the broader premise that antimicrobial function can be achieved through diverse sequence, structural, and mechanistic solutions.

## Discussion

The discovery of antimicrobial peptides is fundamentally constrained by the vastness of sequence space and the cost and throughput limitations of experimental screening. Here, we address these challenges by developing an integrated generative–predictive framework that unifies diffusion-based peptide generation with species-specific activity classification, potency regression, and toxicity prediction. This pipeline enables systematic exploration of peptide design space, producing candidates that preserve key features of natural AMPs while extending into previously underexplored sequence–property regimes.

A central advance of this work is the explicit incorporation of terminal modifications into both computational modeling and experimental validation. Although N- and C-terminal capping is routinely used to improve peptide stability and bioactivity, it is rarely integrated into large-scale computational discovery pipelines. By training predictive models on datasets annotated with terminal chemistry and systematically evaluating capped and uncapped variants experimentally, we demonstrate that terminal modifications exert sequence-dependent but predictable effects on antimicrobial potency, membrane interactions, and toxicity. Across multiple backbones, model-predicted directional changes in activity upon acetylation or amidation were frequently concordant with measured MICs, membrane permeabilization, and depolarization assays. These results support the view that chemical tailoring can be integrated into the discovery workflow rather than relegated to post hoc optimization

The framework’s predictive capacity was further supported by broad species-level modeling and validation. Predictive models were trained across 15 bacterial and fungal species and experimentally evaluated using 120 synthesized peptides tested against 11 clinically relevant pathogens, including ESKAPEE members. This scale of orthogonal validation exceeds that of most AMP discovery studies and demonstrates that predictive enrichment is maintained across diverse taxa rather than being confined to a narrow strain set. The synthesized peptides exhibited elevated cationicity and altered hydrophobic balance relative to database AMPs, consistent with exploration of novel physicochemical space, while toxicity filtering effectively reduced cytotoxic liabilities prior to synthesis.

Mechanistic and structural analyses further highlight the diversity of functional solutions accessed by this pipeline. CD spectroscopy in SDS micelles revealed that the peptides span a wide range of secondary structure propensities, including highly helical, β-enriched, mixed, and largely disordered states, with no global convergence toward α-helicity. Consistent with this structural heterogeneity, permeabilization (NPN uptake) and depolarization (DiSC_3_-5) assays revealed multiple membrane interaction phenotypes, including peptides that strongly permeabilize and depolarize membranes, peptides that decouple these processes, and peptides with limited membrane disruption yet retained antibacterial activity. Importantly, terminal modifications modulated these behaviors in a context-dependent manner without enforcing a single mechanistic pathway.

Finally, selected peptides demonstrated *in vivo* efficacy in a murine skin abscess model of *A. baumannii*, achieving significant early reductions in bacterial burden comparable to polymyxin B. A matched pair of capped and uncapped peptides with identical backbones showed comparable *in vivo* efficacy despite distinct toxicity and membrane-interaction profiles *in vitro*, underscoring the utility of terminal modification for tuning selectivity without compromising antibacterial performance. Together, these results demonstrate that peptides prioritized by the pipeline can translate from computational design to *in vivo* activity.

In sum, this study establishes a scalable, modification-aware paradigm for antimicrobial peptide discovery that integrates generative modeling, predictive filtering, and extensive experimental validation. By explicitly accounting for terminal chemistry, embracing structural and mechanistic diversity, and validating across multiple pathogens and biological scales, this framework advances AMP discovery toward a data-driven strategy for developing next-generation peptide therapeutics to combat multidrug-resistant infections. Nevertheless, some limitations should be considered. The training datasets were compiled from existing AMP databases, which may be biased toward specific peptide classes, species, or assay conditions; as a result, the models may not fully capture the diversity of natural AMPs. In addition, although the diffusion model generates peptides with AMP-like characteristics, generation is constrained by the training distribution and may preferentially produce variations on known motifs rather than entirely new sequence families. The predictive models also relied primarily on SMILES-based representations and physicochemical descriptors, which do not explicitly encode higher-order structural information or dynamic interactions with bacterial membranes that can shape activity and selectivity. Finally, our evaluation of terminal modifications was restricted to two common alterations, N-terminal acetylation and C-terminal amidation. Other biologically relevant chemical modifications such as lipidation, glycosylation, or the incorporation of non-natural amino acids were not explored and may further improve stability, potency, and selectivity. Together, these limitations highlight directions for future methodological and experimental work to enable comprehensive validation and successful clinical translation.

## Methods

### Dataset preparation for generative and filtering models

For the generative model, we compiled a comprehensive AMP dataset by integrating sequences from multiple open-source repositories, including APD ^14–17^, dbAMP ^18–20^, DRAMP ^21–24^, dbaasp ^25–27^, CAMP ^28–30^, LAMP ^31,32^, ParaPep ^33^, phytAMP ^34^, AVPdb ^35^, CancerPPD ^36,67^, AntiTbPdb ^37^, Hemolytik ^38,68^, and milkAMP ^39^, as well as the training datasets of 24 AMP classifiers, including iAMPCN ^40^, ADAM ^41^, iAMP-2L ^42^, AMPfun ^43^, iAMP-CA2L ^44^, MLACP ^45^, AntiCP 2.0 ^46^, ACPred ^47^, ACPred-FL ^48^, ACPred-Fuse ^49^, ACP-DL ^50^, iACP-DRLF ^51^, mACPpred ^52^, AntiFP ^53^, AVPpred ^54^, AVPIden ^55^, AntiTbPred ^56^, AtbPpred ^57^, BIOFIN ^58^, BIPEP ^59^, dPABBs ^60^, HemoPI ^61^, HLPpred-Fuse ^62^, iAntiTB ^63^, AnOxPePred ^64^, and HAPPENN ^65^. After removing sequences containing non-standard residues (B, J, O, U, X, Z) and filtering by length (5–30 amino acid residues), a total of 32,095 AMP sequences were retained as the training set for peptide generation.

For predictive modeling, we curated species-specific activity datasets from dbaasp ^25–27^, yielding 37,119 peptide–species activity pairs across 15 pathogens, including 20,803 active cases (MIC ≤ 32 µg/ml) and 16,316 inactive cases (MIC > 32 µg/ml). Because a single peptide can have MIC measurements against multiple species, these counts refer to peptide–species pairs rather than unique sequences. The largest subsets corresponded to *E. coli* (4,315 active, 3,717 inactive), *S. aureus* (3,742 active, 3,604 inactive), and *P. aeruginosa* (2,990 active, 2,544 inactive). Additional Gram-negative species included *K. pneumoniae* (1,186 active, 815 inactive), *S. enterica* (933 active, 727 inactive), *A. baumannii* (908 active, 573 inactive), and *S. typhimurium* (571 active, 225 inactive). Gram-positive datasets comprised *B. subtilis* (1,720 active, 819 inactive), *S. epidermidis* (1,432 active, 694 inactive), *E. faecalis* (880 active, 656 inactive), *M. luteus* (557 active, 328 inactive), *E. faecium* (314 active, 228 inactive), *L. monocytogenes* (328 active, 184 inactive), and B. cereus (338 active, 221 inactive). For fungal pathogens, *C. albicans* was represented by 1,389 active and 1,011 inactive samples. A complete summary of sample counts is provided in **Table S11**.

For regression modeling, the same DBAASP-derived datasets were used, but raw MIC values (µg/ml) were retained instead of being binarized. When multiple MIC measurements were available for a given peptide–species pair, their average was taken. To normalize the distribution and reduce the impact of extreme outliers, MIC values were transformed as log(MIC + 1), with values above 127 µg/ml capped at 127 prior to transformation. These processed values were then used as regression targets for model training.

For toxicity prediction, we assembled 8,834 peptide entries with haemolytic or cytotoxicity data from DBAASP. Peptides showing <50% haemolysis or <50% cell death at ≥100 µg/ml were labeled as low toxicity (2394 samples), while those exceeding this threshold were considered toxic (6440 samples) (**Table S12**).

### Peptide encoding

We used different methods in different models to encode peptides. For AMP generation, amino-acid sequences could be transformed into matrix representations to use as input data. We treated sequences as test information with each animo acids as tokens. For the AMP prediction, we transformed the amino-acid sequences with terminus modifications into its simplified molecular-input line-entry system (SMILES) format to represent the chemical molecules. The chemical molecules will be treated as graphs which include the edge and atom information. The graph would be as the input for the Chemprop models. We also calculated some physiochemical information as input to represent the molecules, including Morgan fingerprints and rdkit descriptors.

### Diffusion-based generative model for peptide design

For latent diffusion–based peptide generation, we adopted a denoising framework that directly operates on the latent embedding space of ESM-2. Peptide sequences were first tokenized and mapped into embeddings x E R^Zxd^using the ESM-2 encoder. Gaussian noise was iteratively added during the forward process, and a denoiser was trained to reconstruct the corrupted representations in the reverse process by minimizing the mean-squared error between predicted and original embeddings. The denoiser was implemented by repurposing the pretrained ESM-2 transformer, allowing it to directly process continuous latent representations. At each diffusion step, the model was conditioned on the diffusion timestep, and continuous physicochemical properties (net charge and hydrophobic ratio), when provided, were incorporated through a lightweight additive conditioning mechanism. Conditional dropout was applied during training to enable classifier-free guidance at inference, allowing the same model to support both unconditional and conditional generation. The outputs from the ESM-2 transformer layers were further processed by a lightweight multilayer perceptron (MLP) to predict the denoised embeddings. Final sequences were decoded via the ESM-2 language model head by selecting the most probable amino acid at each position, ensuring that generated peptides remained aligned with the pre-trained ESM vocabulary.

Compared with AMP-diffusion, which employs a dedicated denoising transformer that reuses the pretrained ESM2 transformer layers and injects timestep information via sinusoidal embeddings with feature-wise scale–shift modulation, our model directly repurposes the pretrained ESM forward dynamics with minimal architectural changes. This design retains closer alignment with the native ESM representation space while supporting optional continuous property conditioning.

Based on the generative model, we generated 1,000 peptides for each length category (5-30 amino acids). To evaluate generative performance, we randomly selected 10,000 peptides produced by our model and compared their distribution with an equal number of peptides generated by other state-of-the-art approaches, including AMPdesigner, ampdiffusion, ampgan, Apex_duo, basic, BroadAMPGPT, DeepAMP, DiffAMP, Hydra, PepCVAE, and PepDiffusion. The t-SNE embeddings are shown in **Figure S11**, and eight physicochemical properties computed using the modlamp Global Analysis function are presented in **Figure S12A–S12H**. Across methods, subsets of generated peptides were consistently located within the AMP region in physicochemical profiles similar to those of known AMPs. the t-SNE space. The physicochemical profiles are also broadly similar to those of known AMPs. To benchmark generative quality, we applied sequential and alternative filtering pipelines (**Figures S12I–S12K**). In the multi-step pipeline, ours consistently achieved the highest success rate, retaining a substantially larger fraction of candidates at each stage relative to other approaches. Application of the APEX-based potency filter (average median predicted MIC <80 μM) further confirmed this advantage, as ours yielded the greatest proportion of retained peptides both before (**Figure S12J**) and after (**Figure S12K**) redundancy reduction with CD-HIT at 70% sequence identity. Although success rates declined across all methods after redundancy removal, the relative advantage of ours was maintained, underscoring its superior capacity to generate potent and novel AMP candidates.

### Classification models for predicting peptide antibacterial activity

To predict peptide antibacterial activity, we implemented Chemprop, a software package based on directed message-passing neural networks (D-MPNNs). Each peptide sequence was converted into its SMILES representation and subsequently transformed into a molecular graph, with atoms and bonds represented as nodes and edges, respectively, in order to incorporate terminal modifications. Iterative graph convolutional operations were performed to aggregate local atomic and bond information, resulting in a fixed-length molecular embedding.

To assess the effect of molecular features on predictive performance, we benchmarked four model variants: (1) Chemprop with no additional features, (2) Chemprop augmented with RDKit-computed physicochemical descriptors, (3) Chemprop with Morgan fingerprint features, and (4) Chemprop with Morgan count fingerprint features. In each case, the molecular embeddings generated by the D-MPNNs were concatenated with the selected feature set (if provided) before being passed to a fully connected feed-forward neural network for classification. The model output was a probability score between 0 and 1, representing the likelihood that the peptide was active. Training data were derived from curated peptide–MIC pairs from dbaasp database. For peptides with multiple MIC values against the same bacterial species, the average MIC was used. Activity labels were binarized using a threshold of 32 µg/ml (≤32 µg/ml: active; >32 µg/ml: inactive). Species-specific models were trained by splitting the data into training and test sets (80:20 ratio). Hyperparameter optimization was performed using a limited grid search over learning rate, hidden layer size, message-passing steps, and dropout, as summarized in **Table S13**. The best-performing configuration was selected for each Chemprop variant. On the training sets, 10-fold cross-validation was performed, and ensemble predictions were generated by averaging across the ten folds. Performance comparisons across the four model variants revealed that the Morgan count fingerprints consistently improved predictive accuracy relative to other feature sets. As a result, the Morgan count–augmented Chemprop model was selected as the primary classifier in the funnel selection pipeline.

### Regression models for predicting peptide antibacterial activity

In addition to binary classification, we implemented a regression stage to predict quantitative MIC values (measured in µg/ml) for each peptide–species pair. For this task, multiple MIC measurements from DBAASP were averaged and then transformed using log(MIC + 1) to normalize the distribution. To limit the effect of extreme outliers, MIC values greater than 127 µg/ml were capped at this threshold before transformation. Species-specific Chemprop models were trained using SMILES-derived molecular graphs, optionally augmented with RDKit physicochemical descriptors, Morgan fingerprints, or Morgan count fingerprints. The models were optimized using an L2-loss objective. Data were randomly split into training and test sets (80:20 ratio), and 10-fold cross-validation was applied on the training data. Hyperparameter optimization was performed using a limited grid search over learning rate, hidden layer size, message-passing steps, and dropout, as summarized in **Table S14**. The best-performing configuration was selected for each Chemprop variant. Ensemble predictions were obtained by averaging the outputs of ten models. Regression performance was assessed by computing the Pearson correlation coefficients and Spearman rank correlation coefficient between predicted and experimentally measured MIC values. Consistent with the classification stage, regression models incorporating Morgan count features achieved the best predictive performance across multiple species.

### Toxicity prediction models

Model training followed the same Chemprop framework as described for antibacterial classification, with peptides represented by SMILES-derived molecular graphs and optionally augmented with RDKit descriptors, Morgan fingerprints, or Morgan count fingerprints. Data were split 80:20 into training and test sets, and 10-fold cross-validation was applied to the training portion. Hyperparameter optimization was performed using a limited grid search over learning rate, hidden layer size, message-passing steps, and dropout, as summarized in **Table S15**. The best-performing configuration was selected for each Chemprop variant. Model performance was assessed using the area under the ROC curve (AUC), and Morgan count–augmented models consistently achieved the highest AUC values. These were therefore adopted for filtering potentially haemolytic or cytotoxic peptides in the funnel selection pipeline.

### Down-selection of generated peptides for synthesis and testing

All generated sequences were expanded by applying two common terminal modifications, N-terminal acetylation and C-terminal amidation, resulting in four peptide variants for each original sequence (unmodified, N-acetylated, C-amidated, and N-acetylated/C-amidated). Each variant was then evaluated by the predictive models, and peptides were retained only if they satisfied the following thresholds for each bacterial species: (i) predicted antibacterial classification probability ≥ 0.8, (ii) predicted MIC regression output corresponding to log(MIC + 1) < 5, and (iii) predicted toxicity probability < 0.2. Following this model-based filtering, to ensure novelty relative to existing data, all unmodified backbone sequences were first compared against the training dataset, and sequences sharing ≥70% identity were removed using CD-HIT. This step eliminated candidates that were too similar to known peptides. To preserve diversity within the candidate pool itself, the remaining unmodified sequences were clustered again using CD-HIT at 70% identity, and only representative sequences from each cluster were retained. For each bacterial species, the retained peptides were subsequently ranked in descending order based on a composite prediction score for antibacterial activity defined as:

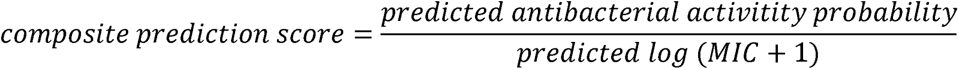

Top-ranked peptides from each species were shortlisted. Since experimental validation in our laboratory was restricted to 11 bacterial species, only these were considered in the final candidate selection: *A. baumannii* (ATCC 19606), *E. coli* (ATCC 11775), *K. pneumoniae* (ATCC 13883), *P. aeruginosa* (PAO1), *S. enterica* (ATCC 9150), *S. Typhimurium* (ATCC 700720), *B. subtilis* (ATCC 23857), *S. aureus* (ATCC 12600), *L. monocytogenes* (ATCC 19111), *E. faecalis* (ATCC 700802, VRE), and *E. faecium* (ATCC 700221, VRE). The species-specific lists were merged, and peptides were further ranked according to the number of species for which they achieved high scores. Candidates with the broadest coverage across species were prioritized for synthesis and experimental validation. The peptides that passed each stage are summarized in **Data S2**. The experimental analysis was conducted using µM as the activity unit. However, model training and testing were performed using µg/mL, as the DBAASP-derived dataset reports activity values in µg/mL. For comparison with the experimental results, the predicted values were converted from µg/mL to µM based on the molecular weight of each peptide.

### Comparative sequence analysis of generated peptides

Generated peptide sequences were compared against natural proteins using DIAMOND (blastp mode; v2.1.10) with the --more-sensitive setting, an e-value threshold of 1e-1, and no limit on reported alignments, against the UniProt reference database (Release 2025_03). DIAMOND outputs (tabular .m8 format) were processed to calculate query coverage, and hits were retained if they satisfied e-value ≤ 1e-1, sequence identity ≥ 20%, and query coverage ≥ 0.7. For each query peptide, the top five hits ranked by bit score were retained.

Filtered hits were annotated in UniProt to obtain protein and species information. Aligned fragments corresponding to the matched regions were extracted from the UniProt reference sequences for downstream comparative analyses. Species distributions of the matched proteins were summarized, and phylogenetic trees were visualized and annotated using the Interactive Tree of Life (iTOL) web tool to highlight the taxonomic context of the alignments.

### Peptide sequence analysis

Physicochemical properties of the peptides, including net charge, charge density, isoelectric point (pI), instability index, aromaticity, aliphatic index, Boman index, and hydrophobic residues content, were calculated using the GlobalDescriptor module implemented in the modlamp Python package. The instability index is based on dipeptide composition, aromaticity is defined as the fraction of phenylalanine (F), tryptophan (W), and tyrosine (Y), and hydrophobic ratio represents the proportion of alanine (A), cysteine (C), phenylalanine (F), isoleucine (I), leucine (L), methionine (M), and valine (V) residues.

### Peptide Synthesis

All peptides used in the experiments were purchased from AAPPTec and synthesized by solid-phase peptide synthesis using the Fmoc strategy.

### Culturing conditions and bacterial strains

In this study, we used the following pathogenic bacterial strains obtained from the American Type Culture Collection (ATCC): *Acinetobacter baumannii* ATCC 19606, *Escherichia coli* ATCC 11775, *Klebsiella pneumoniae* ATCC 13883, *Pseudomonas aeruginosa* PAO1, *Salmonella enterica* ATCC 9150, *Salmonella enterica* Typhimurium ATCC 700720, *Bacillus subtilis* ATCC 23857, *Staphylococcus aureus* ATCC 12600, *Listeria monocytogenes* ATCC 19111, *Enterococcus faecalis* ATCC 700802 (vancomycin-resistant strain), and *Enterococcus faecium* ATCC 700221 (vancomycin-resistant strain). Pseudomonas Isolation (*Pseudomonas aeruginosa* strains) agar plates were exclusively used in the case of *Pseudomonas* species. All the other pathogens were grown in Luria-Bertani (LB) broth and on LB agar. In all the experiments, bacteria were inoculated from one-isolated colony and grown overnight (16 h) in liquid medium at 37 °C. In the following day, inoculums were diluted 1:100 in fresh media and incubated at 37 °C to mid-logarithmic phase.

### Human cells and serum

Human embryonic kidney (HEK293T) cells were obtained from American Type Culture Collection (CRL-3216). Red blood cells (RBCs) and human serum were purchased from Zen-Bio. The RBC samples were obtained from the same certified healthy donor (blood type A^−^).

### Minimal inhibitory concentration determination

Broth microdilution assays were performed to determine the minimum inhibitory concentration (MIC) values of each peptide. Peptides were added to nontreated polystyrene microtiter 96-well plates and 2-fold serially diluted in sterile water from 1 to 64 μmol L^−1^. Bacterial inoculum at 4×10^6^ CFU mL^−1^ in LB medium was mixed 1:1 with the peptide. The MIC was defined as the lowest concentration of peptide able to completely inhibit the bacterial growth after 24 h of incubation at 37 °C. All assays were done in three independent replicates.

### Circular dichroism experiments

The circular dichroism experiments were conducted using a J1500 circular dichroism spectropolarimeter (Jasco) in the Biological Chemistry Resource Center (BCRC) at the University of Pennsylvania. Experiments were performed at 25 °C, the spectra graphed are an average of three accumulations obtained with a quartz cuvette with an optical path length of 1.0 mm, ranging from 260 to 190 nm at a rate of 50 nm min^−1^ and a bandwidth of 0.5 nm. The concentration of all peptides tested was 50 μmol L^−1^, and the measurements were performed in sodium dodecyl sulfate (SDS) in water at 10 mmol L^−1^, with baseline recorded prior to measurement. A Fourier transform filter was applied to minimize background effects. Secondary structure fraction values were calculated using the single spectra analysis tool on the server BeStSel^19^.

### Outer membrane permeabilization assays

N-phenyl-1-napthylamine (NPN) uptake assay was used to evaluate the ability of the peptides to permeabilize the bacterial outer membrane. Inocula of *A. baumannii* ATCC 19606 were grown to an OD at 600 nm of 0.4 mL^−1^, centrifuged (9,391 ×g at 4 °C for 10 min), washed and resuspended in 5 mmol L^−1^ HEPES buffer (pH 7.4) containing 5 mmol L^−1^ glucose. The bacterial solution was added to a white 96-well plate (100 μL per well) together with 4 μL of NPN at 0.5 mmol L^−1^. Consequently, peptides diluted in water were added to each well, and the fluorescence was measured at λ_ex_ = 350 nm and λ_em_ = 420 nm over time for 45 min. The relative fluorescence was calculated using the untreated control (buffer + bacteria + fluorescent dye) and polymyxin B (positive control) as baselines and the following equation was applied to reflect % of difference between the baselines and the sample:

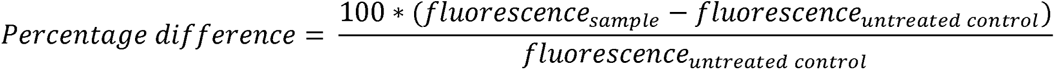

### Cytoplasmic membrane depolarization assays

The cytoplasmic membrane depolarization assay was performed using the membrane potential-sensitive dye 3,3’-dipropylthiadicarbocyanine iodide (DiSC3-5). *A. baumannii* ATCC 19606 and *P. aeruginosa* PAO1 in the mid-logarithmic phase were washed and resuspended at 0.05 OD mL^−1^ (optical value at 600 nm) in HEPES buffer (pH 7.2) containing 20 mmol L^−1^ glucose and 0.1 mol L^−1^ KCl. DiSC3-5 at 20 μmol L^−1^ was added to the bacterial suspension (100 μL per well) for 15 min to stabilize the fluorescence which indicates the incorporation of the dye into the bacterial membrane, and then the peptides were mixed 1:1 with the bacteria to a final concentration corresponding to their MIC100 values. Membrane depolarization was then followed by reading changes in the fluorescence (λ_ex_ = 622 nm, λ_em_ = 670 nm) over time for 60 min. The relative fluorescence was calculated using the untreated control (buffer + bacteria + fluorescent dye) and polymyxin B (positive control) as baselines and the following equation was applied to reflect % of difference between the baselines and the sample:

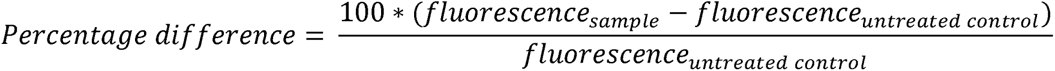

### Hemolytic activity assays

To evaluate the release of hemoglobin from human erythrocytes upon treatment of each of the encrypted peptides, human red blood cells (RBCs) were obtained from ZenBio (male donor, blood type A^−^) obtained from heparin anti-coagulated blood. RBCs were washed with PBS (pH 7.4) four times by centrifugation at 800 ×g for 10 min. Aliquots of 200-fold diluted cells (75 μL) were mixed with peptide solution (0.78-100 μmol L^−1^; 75 μL), and the mixture was incubated for 4 h at room temperature. After the incubation, the plate was centrifuged at 1,300 ×g for 10 min to precipitate cells and debris, and 100 μL of supernatant from each well were transferred to a new 96-well plate for absorbance reading (405 nm) using an automatic plate reader. The percentage of hemolysis was defined by comparison with negative control (samples containing PBS) and positive control [samples containing 1% (v/v) SDS in PBS solution].

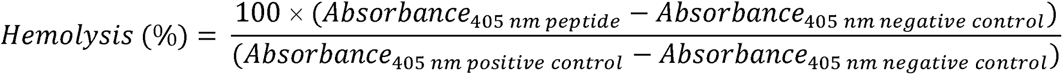

### Cytotoxicity assays

The cells were cultured in high-glucose Dulbecco’s modified Eagle’s medium supplemented with 1% penicillin and streptomycin (antibiotics) and 10% fetal bovine serum and grown at 37□°C in a humidified atmosphere containing 5% CO_2_.

One day prior to the experiment, 100□μL aliquots of human embryonic kidney (HEK293T) cells, at a concentration of 50,000□cells per mL, were seeded into each well of 96-well plates (5,000 cells per well). Following cell attachment, the HEK293T cells were treated with increasing concentrations of peptides (ranging from 8 to 128□μmol[L^−1^) and incubated for 24 h. After the exposure period, cytotoxicity was assessed using the 3-(4,5-dimethylthiazol-2-yl)-2,5-diphenyltetrazolium bromide (MTT) assay. Specifically, the MTT reagent was prepared at a concentration of 0.5□mg[mL^−1^ in phenol red-free medium and used to replace the peptide-containing supernatants (100□μL per well). The plates were then incubated for 4□h at 37□°C in a humidified atmosphere with 5% CO□, facilitating the formation of insoluble formazan crystals. These crystals were subsequently dissolved in 0.04□mol□L^−1^ hydrochloric acid prepared in anhydrous isopropanol. Absorbance was measured at 570□nm using a spectrophotometer to quantify cell viability. All experiments were conducted in triplicate (three biological replicates).

### Skin abscess infection mouse model

The back of six-week-old female CD-1 mice under anesthesia were shaved and injured with a superficial linear skin abrasion made with a needle. An aliquot of *A. baumannii* ATCC 19606 (7.66×10^5^ CFU mL^−1^; 20 μL) previously grown in LB medium until 0.5 OD mL^−1^ (optical value at 600 nm) and then washed twice with sterile PBS (pH 7.4, 9,391 ×g for 3 min) was added to the scratched area. Peptides diluted in sterile water at their MIC value were administered to the wounded area 1 h post-infection. Two- and four-days post-infection, animals were euthanized, and a uniform excision of the scarified skin was excised, homogenized using a bead beater (25 Hz for 20 min), 10-fold serially diluted, and plated on McConkey agar plates for CFU quantification. The experiments were performed using six mice per group. Mice were single-housed to avoid cross-contamination and maintained under a 12-hour light/dark cycle at 22□°C with humidity controlled at 50%. The skin abscess infection mouse model was revised and approved by the University Laboratory Animal Resources (ULAR) from the University of Pennsylvania (Protocol 806763).

### Reproducibility of the experimental assays

All assays were performed in three independent biological replicates as indicated in each figure legend and in the Experimental Models and Methods details sections. The values obtained for hemolytic activity were estimated by non-linear regression based on the screen of peptides in a gradient of concentrations and represent the hemolytic concentration values needed to lyse and kill 50% of the cells present in the experiment. In the skin abscess and thigh infection mouse models, we used six mice per group following established protocols approved by the University Laboratory of Animal Resources (ULAR) of the University of Pennsylvania.

### Quantification and statistical analysis

All statistical analyses were performed in Python and R using the scikit-learn, SciPy, tidyverse, pROC, and ggpubr packages. For classification tasks, metrics reported included accuracy (ACC), precision, recall, F1, and confusion-matrix counts (TP, FP, TN, FN). For regression tasks, performance was assessed using mean squared error (MSE), Pearson correlation, and Spearman correlation in scikit-learn and SciPy. ROC curves and AUC with 95% confidence intervals were computed using the pROC package in R. For physicochemical descriptor comparisons, two-sided Wilcoxon rank-sum tests were applied using ggpubr. The choice of statistical test was based on the data type and experimental design. Unless otherwise specified, all tests were two-sided, and no data were excluded from the analyses. Statistical details can be found in the figure legends.

In the mouse experiments, all the raw data were log_10_ transformed and the statistical significance was determined using one-way ANOVA followed by Dunnett’s test. All the p values are shown for each of the groups, all groups were compared to the untreated control group. All calculation and statistical analyses of the experimental data were conducted using GraphPad Prism v.10.3. Statistical significance between different groups was calculated using the tests indicated in each figure legend. No statistical methods were used to predetermine sample size.

## Data availability

The antimicrobial peptides analyzed in this study were obtained from public available databases, including APD (https://aps.unmc.edu), dbAMP (https://ycclab.cuhk.edu.cn/dbAMP/), DRAMP (https://dramp.cpu-bioinfor.org), dbaasp (https://dbaasp.org/home), CAMP (https://camp.bicnirrh.res.in), LAMP (http://biotechlab.fudan.edu.cn/database/lamp/index.php), ParaPep (https://webs.iiitd.edu.in/raghava/parapep/home.php), phytAMP (https://webs.iiitd.edu.in/raghava/satpdb/catalogs/phytamp/), AVPdb (https://webs.iiitd.edu.in/raghava/satpdb/catalogs/avpdb/), CancerPPD (https://webs.iiitd.edu.in/raghava/cancerppd/cancerppd2.php), AntiTbPdb (https://webs.iiitd.edu.in/raghava/antitbpdb/), Hemolytik (http://crdd.osdd.net/raghava/hemolytik/), and milkAMP (http://milkampdb.org/). The curated datasets used in this study are also available via Mendeley Data (https://data.mendeley.com/preview/5bknbrhyv5?a=fd63ac9e-60a2-4ca5-b0bb-207b9071c3b7). Further information and requests for resources should be directed to Cesar de la Fuente-Nunez.

## Code availability

Termini is available at GitLab (https://gitlab.com/joy1314bubian/capamp.git)

## Acknowledgments

C.F.-N. holds a Presidential Professorship at the University of Pennsylvania and acknowledges funding from the National Institute of General Medical Sciences of the National Institutes of Health under award number R35GM138201 and the Defense Threat Reduction Agency (DTRA; HDTRA1-21-1-0014). J.S. acknowledges funding support from the National Health and Medical Research Council (NHMRC) of Australia (grant nos. APP1127948, APP1144652, APP2036864), and the Major and Seed Inter-Disciplinary Research projects awarded by Monash University. F.L. acknowledges support from the NHMRC Investigator Grant (grant no. 2041439). The South Australian immunoGENomics Cancer Institute (SAiGENCI) is supported by funding from the Australian Government. C.L. acknowledges funding support from an NHMRC Ideas Grant (2024/GNT2037597) and an Australian Research Council Future Fellowship (FT240100798). We thank de la Fuente Lab members for insightful discussions. Figures created with BioRender.com are attributed as such.

## Author contributions

Jing Xu and Jiangning Song designed the computational study. Marcelo D. T. Torres and Cesar de la Fuente-Nunez designed the experimental study. Marcelo D. T. Torres performed the experiments and interpreted the experimental data. Jing Xu performed all computational analyses. Chen Li assisted with the computational analyses. Cesar de la Fuente-Nunez, Jiangning Song, and Fuyi Li supervised the study. Jing Xu and Marcelo D. T. Torres wrote the original draft of the manuscript. All authors reviewed, edited, and approved the final manuscript.

