## Supplementary Information file for "Modification-aware AI enables terminal chemical modifications for peptide design and discovers potent antimicrobials"

**Supplementary Figures S1-S11**

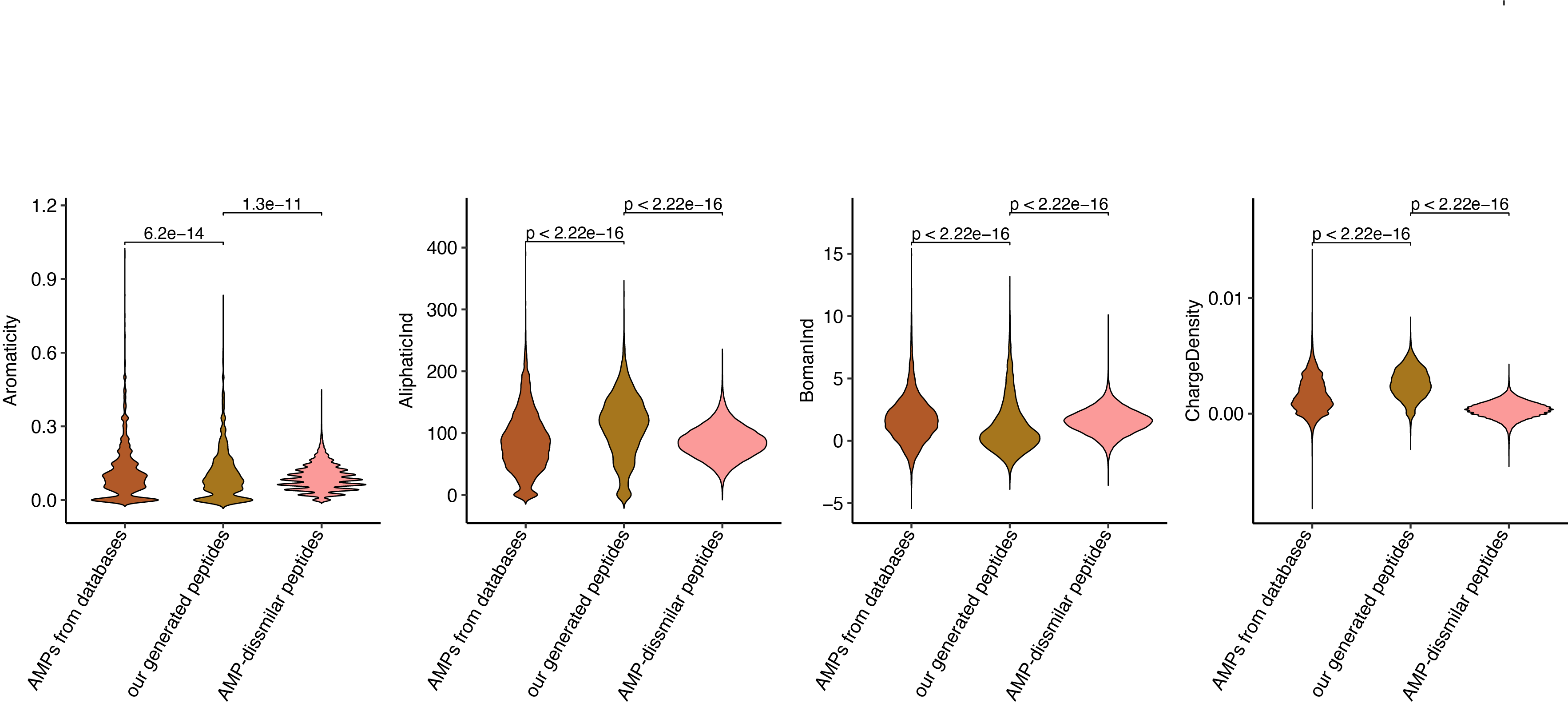

#### **Figure S1. Distribution of physicochemical properties.** Violin plots show aromaticity (defined as the fraction of aromatic residues, phenylalanine (F), tryptophan (W), and tyrosine (Y), in the sequence), aliphatic index, Boman index, and charge density across the three peptide groups.

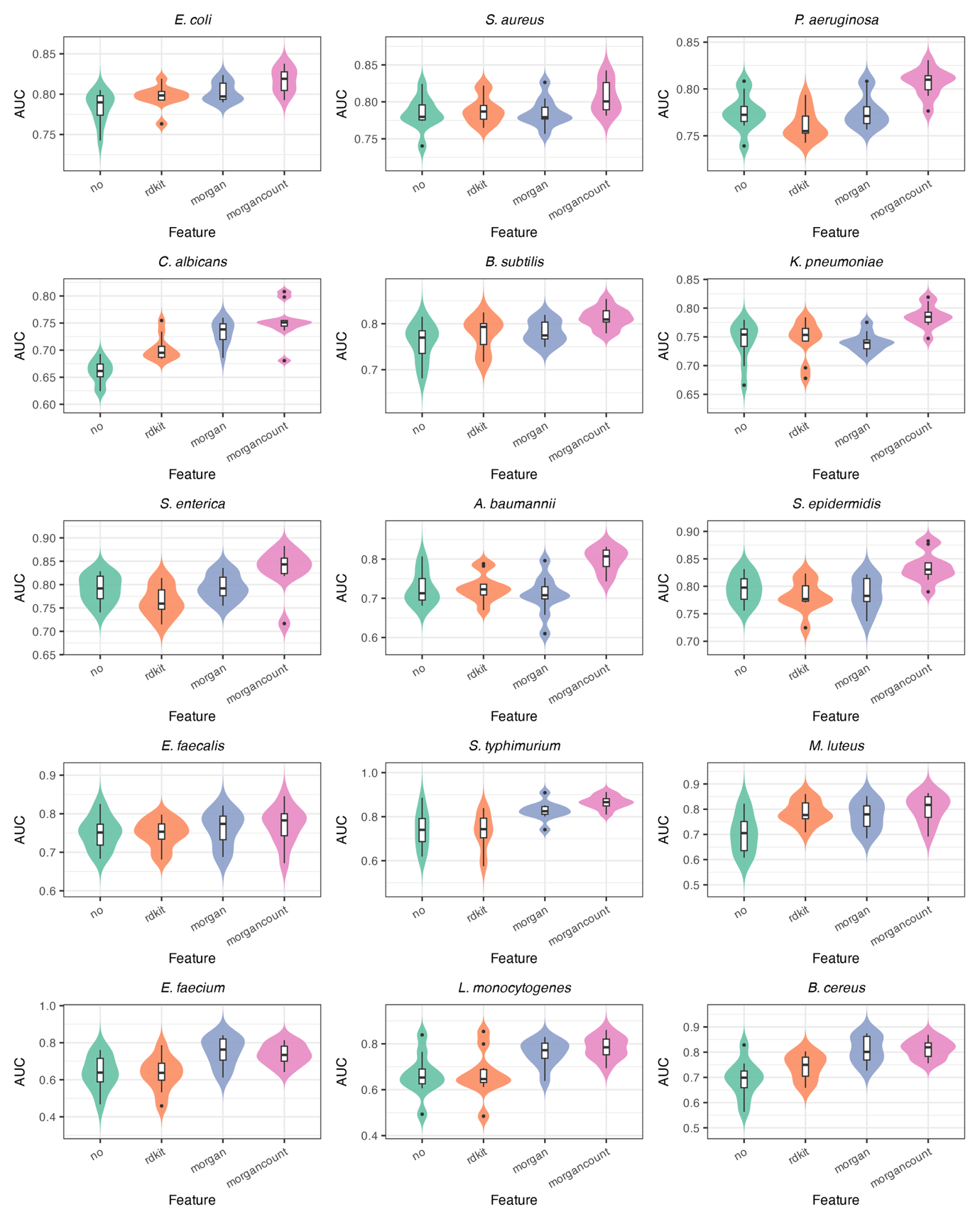

**Figure S2. Performance of classification models evaluated by 10-fold cross-validation.** Violin plots show the distribution of AUC values from 10-fold cross-validation on curated species-level activity datasets. Across species, ensembles of Chemprop classifiers augmented with Morgan fingerprint features consistently achieved the highest predictive performance compared with alternative feature representations.

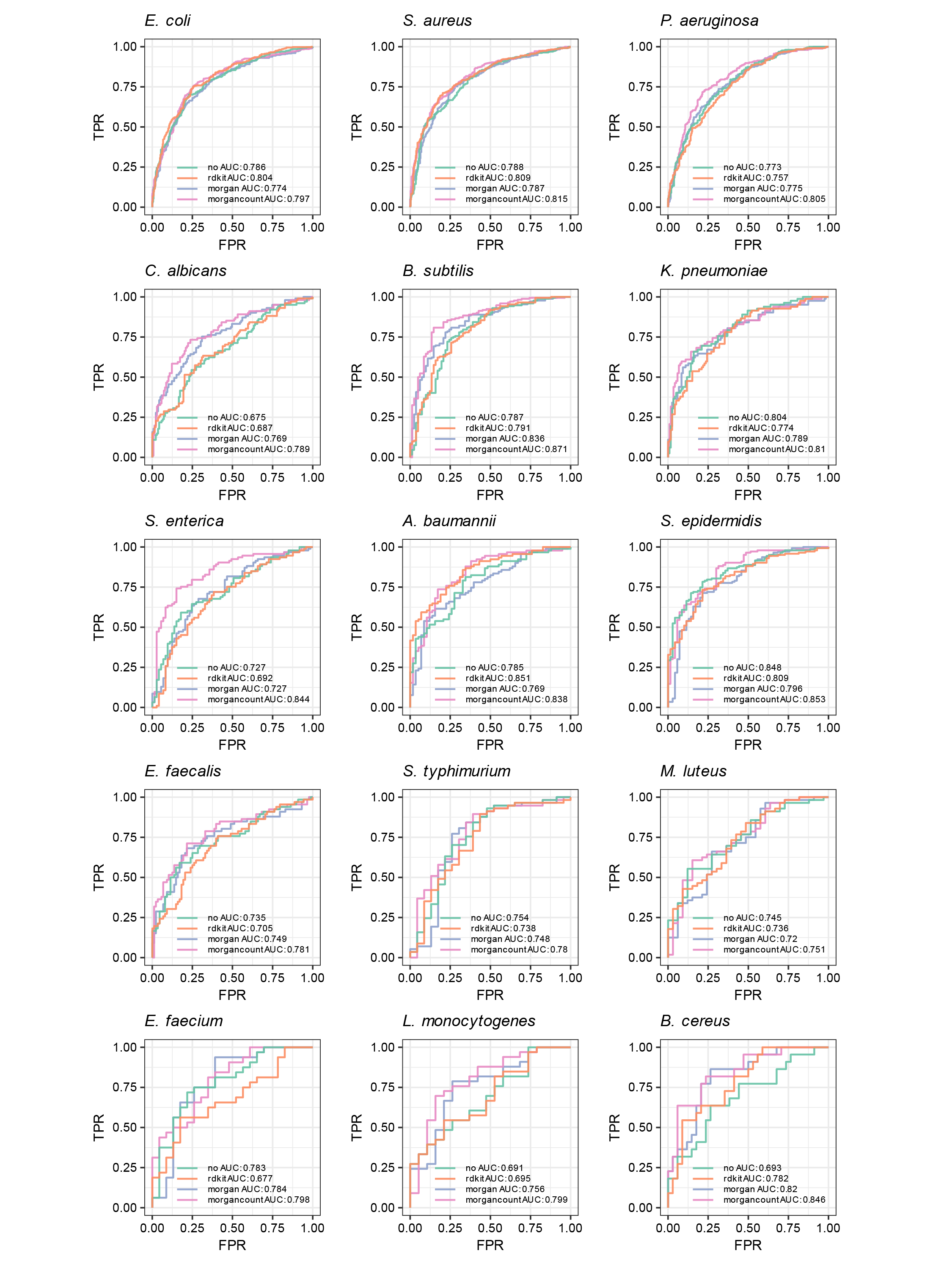

**Figure S3. ROC analysis of classification models on held-out test sets.** Receiver operating characteristic (ROC) curves with corresponding AUC values are shown for models evaluated on 20% held-out test sets from the curated species-level datasets. Ensembles of Chemprop classifiers augmented with Morgan fingerprint features consistently yielded the best predictive performance across species.

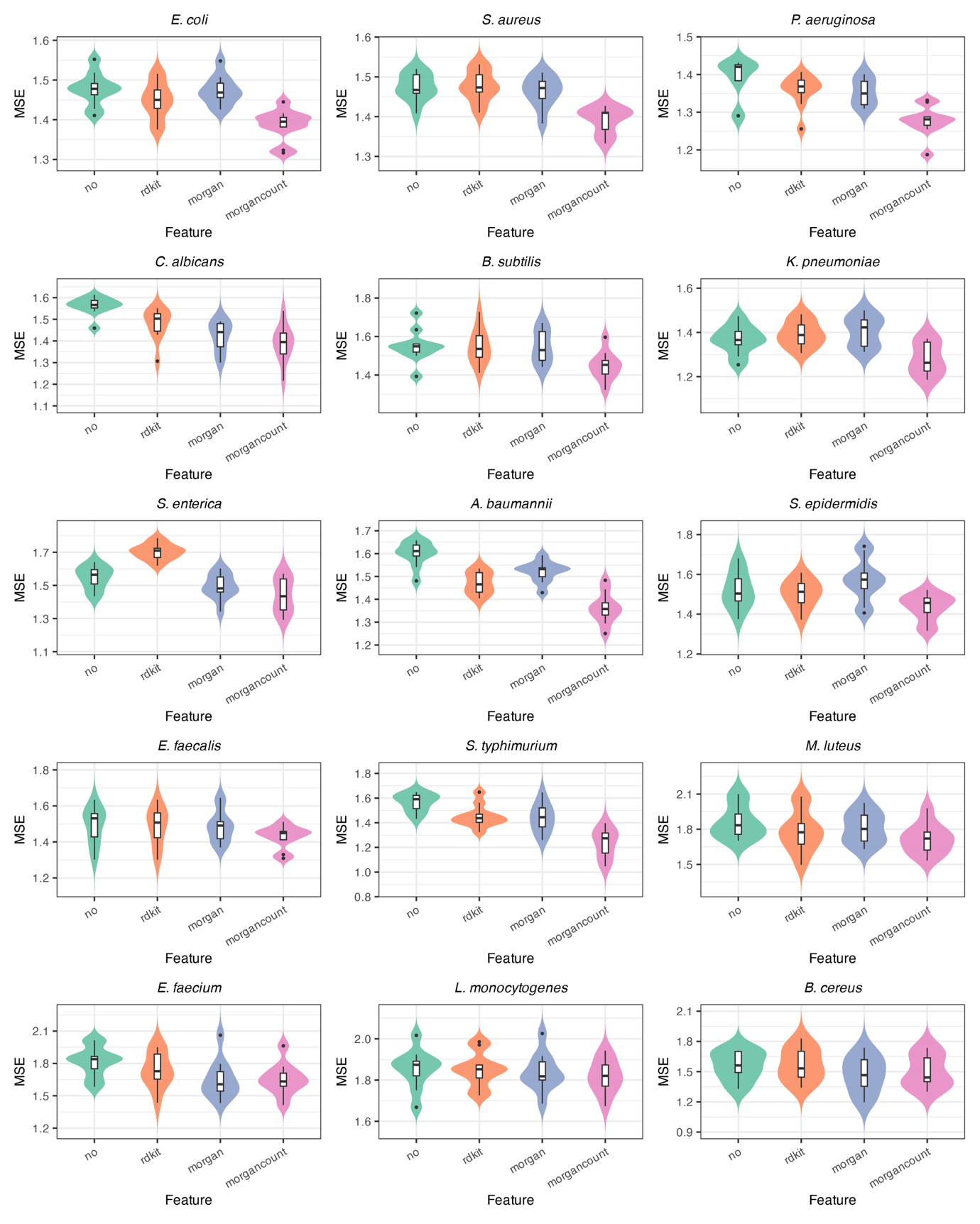

**Figure S4. Performance of regression models evaluated by 10-fold cross-validation.** Violin plots show the distribution of MSE values from 10-fold cross-validation on curated species-level activity datasets.

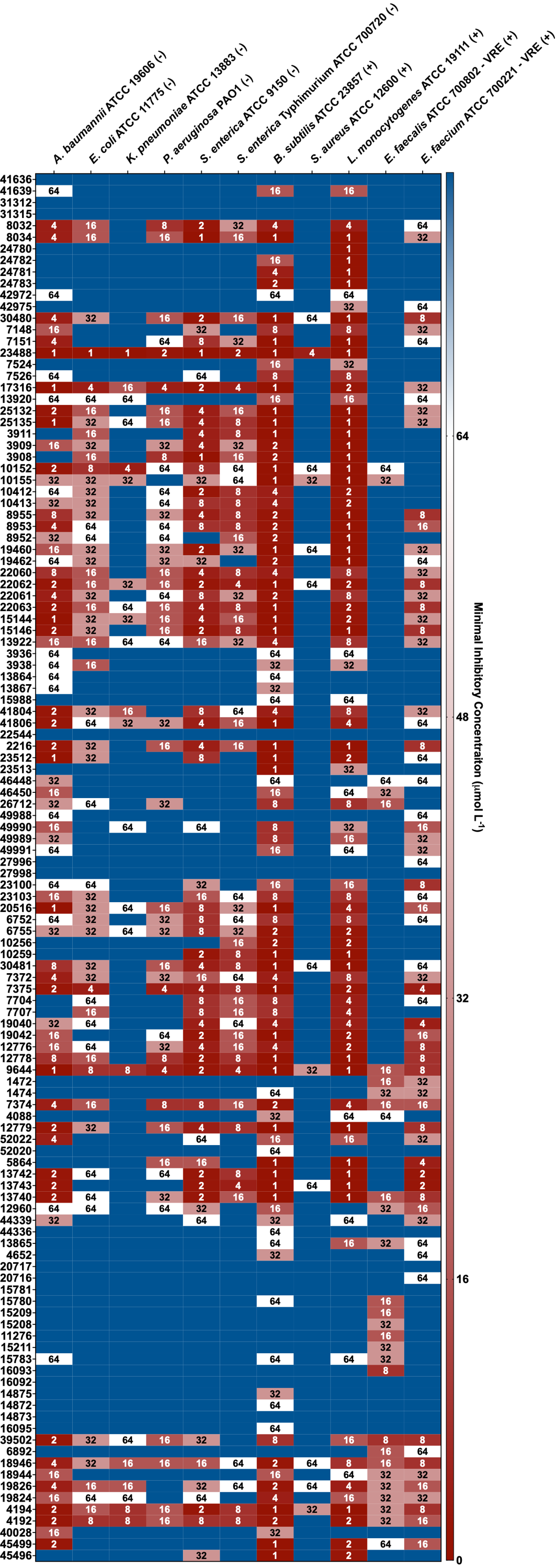

**Figure S5. Antimicrobial activity of tested peptides.** Heat map of minimum inhibitory concentrations (MICs, μmol L^-1^) against 11 clinically relevant Gram-negative (–) and Gram-positive (+) pathogens, including antibiotic-resistant strains. Bacteria (10^5^ CFU) were incubated with serial peptide dilutions (0–64 μmol L^-1^) at 37 °C, and growth was quantified by OD_600_ after 24 h. MIC values represent the mode of replicate measurements for each condition.

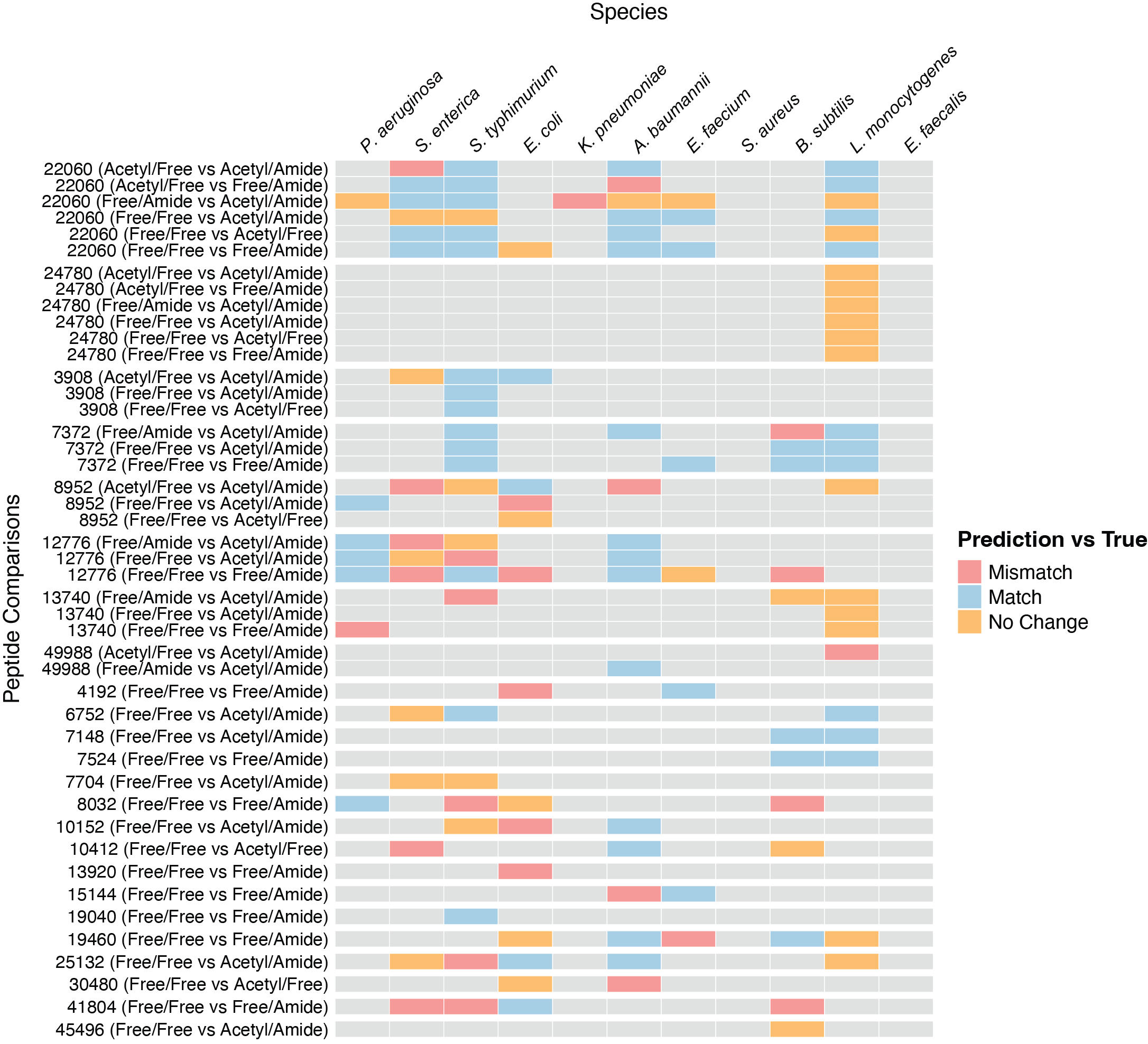

**Figure S6. Directional consistency of predicted and experimental modification effects.** Heatmap of pairwise comparisons of terminally modified peptide variants across bacterial strains. Each row represents a peptide sequence with two different terminal variants, and each column corresponds to a bacterial strain. Colours indicate whether predicted and experimental MIC changes agreed (blue, Match), disagreed (red, Mismatch), or could not be resolved (yellow, Neutral). Neutral denotes cases where experimental MIC values were identical between variants due to the two-fold serial dilution resolution of broth microdilution assays. Grey cells denote cases where the experimental MIC exceeded the highest tested concentration (64 μmol L^-1^), indicating no detectable activity within the assay range. Rows are grouped by peptide sequence and ranked according to the number of pairwise comparisons. Two peptide sequences (22060 and 24780) were selected for complete modification comparisons, providing representative examples of the directional analysis.

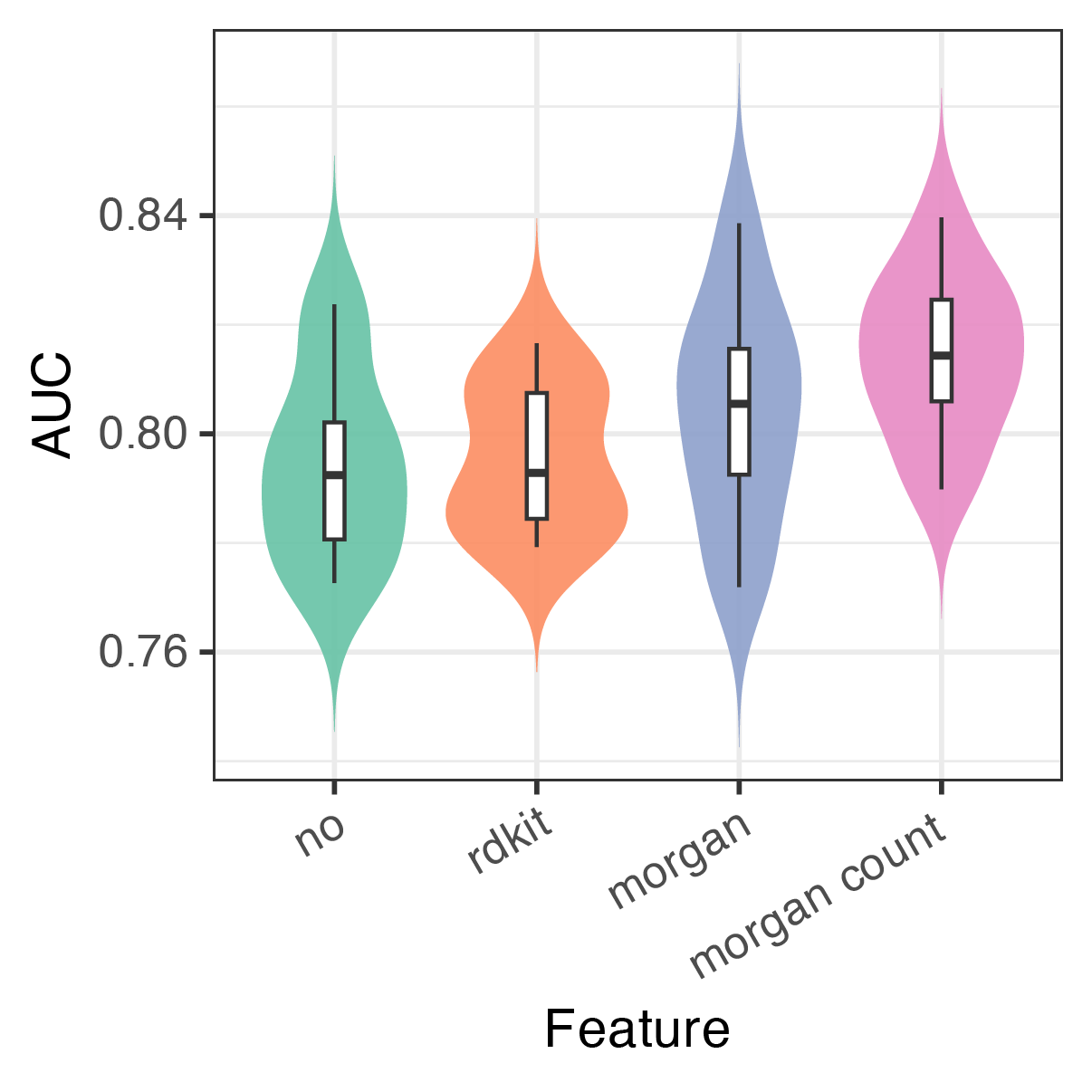

**Figure S7. Performance of toxicity prediction models evaluated by 10-fold cross-validation.** Violin plots show the distribution of AUC values from 10-fold cross-validation on the curated dataset.

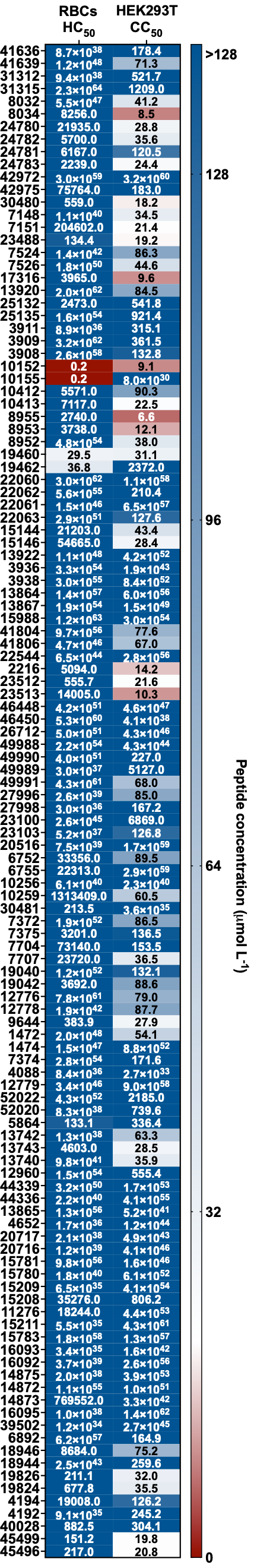

**Figure S8.** **Cytotoxicity and hemolysis of tested peptides.** Hemolytic (HC_50_) and cytotoxic (CC_50_) concentrations, defined as the peptide concentration causing 50% lysis of red blood cells or reduction in HEK293T cell viability, respectively. Values were obtained by nonlinear regression of dose-response curves. Data represent three independent experiments.

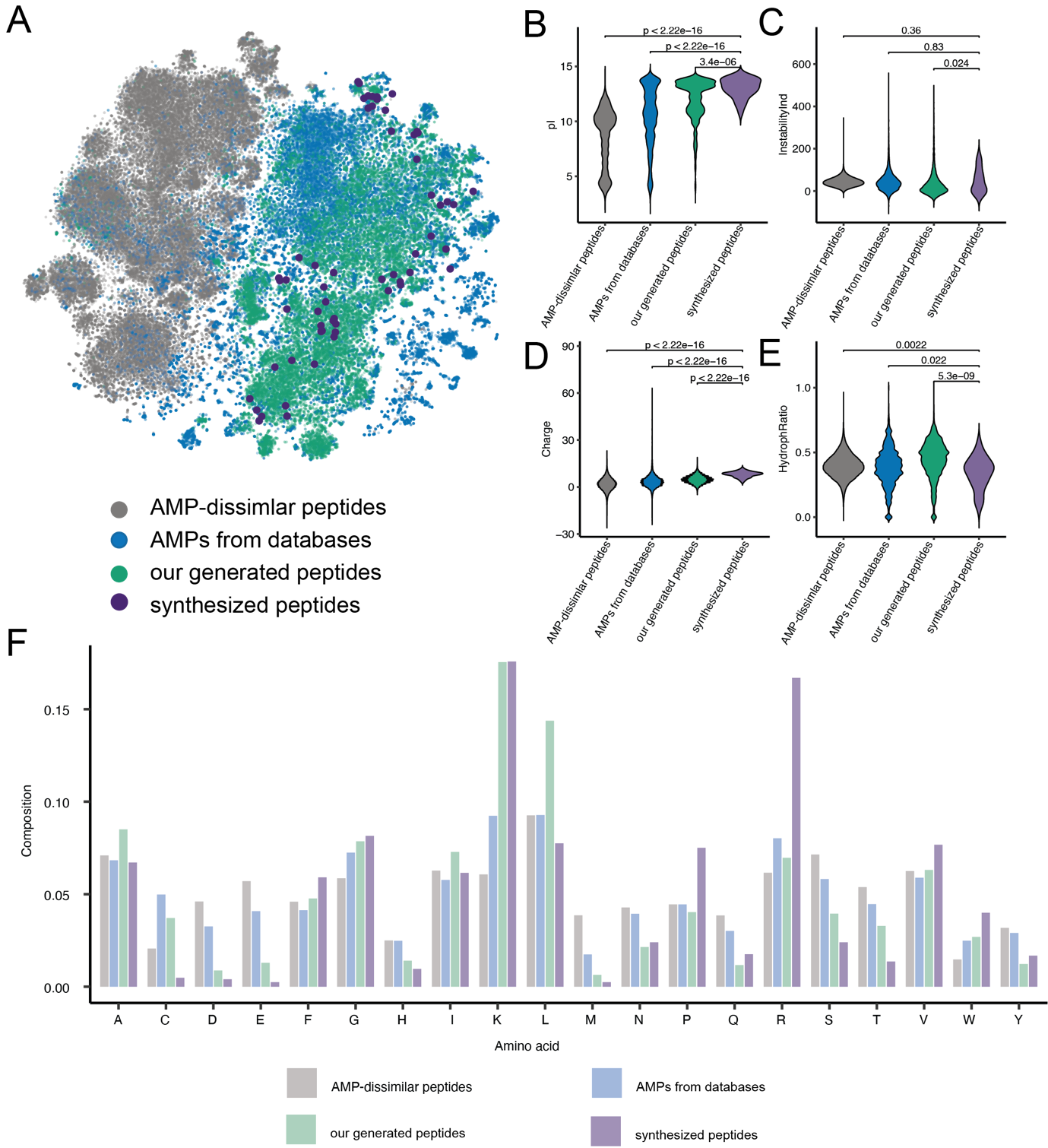

**Figure S9. Physicochemical features distinguish candidate peptides from known AMPs.** **(A)** t-SNE embeddings of peptide sequence space, including AMP-dissimilar peptides (gray), AMPs from databases (blue), our generated peptides (green), and synthesized peptides (purple). The synthesized peptides localized to peripheral or sparsely populated regions, suggesting they occupy distinct sequence space. **(B–E)** Violin plots of key physicochemical parameters, including **(B)** isoelectric point (pI), **(C)** instability index (calculated from dipeptide composition using empirically derived instability weight values), **(D)** net charge, and **(E)** hydrophobic ratio [defined as the proportion of alanine (A), cysteine (C), phenylalanine (F), isoleucine (I), leucine (L), methionine (M), and valine (V) residues over other residues of the sequence]. These physicochemical parameters were calculated using the Global Analysis function in the modlamp Python package. Compared with database AMPs, both generated and synthesized peptides exhibited significantly higher pI and net charge, comparable instability indices, and reduced hydrophobic ratios. **(F)** Amino acid composition across peptide classes, showing enrichment of lysine (K), proline (P), arginine (R), and valine (V), together with reduced methionine (M) in synthesized peptides relative to known AMPs. Together, these analyses indicate that candidate peptides represent a class of sequences with physicochemical features distinct from classical AMPs.

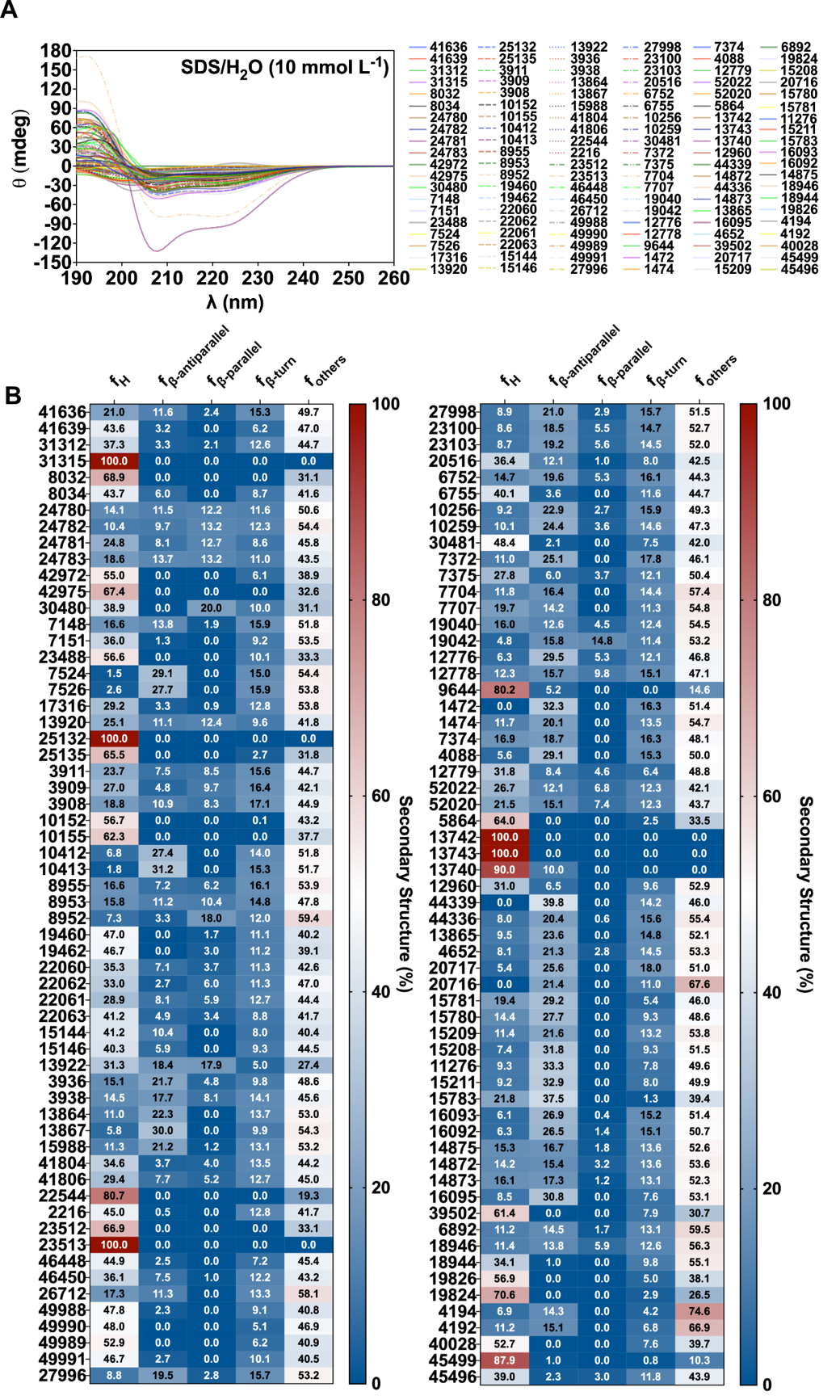

**Figure S10. Secondary structure of tested peptides.** **(A)** Circular dichroism (CD) spectra acquired using a J-1500 Jasco spectropolarimeter. Peptides (50 μmol L^-1^) were analyzed in 10 mmol L^-1^ SDS in water at 25 °C (1 mm path length), recorded from 260-190 nm (bandwidth 0.5 nm, scan rate 50 nm min^-1^; three accumulations). **(B)** Heat map shows the percentage of secondary structure for each peptide calculated using the BeStSel algorithm.

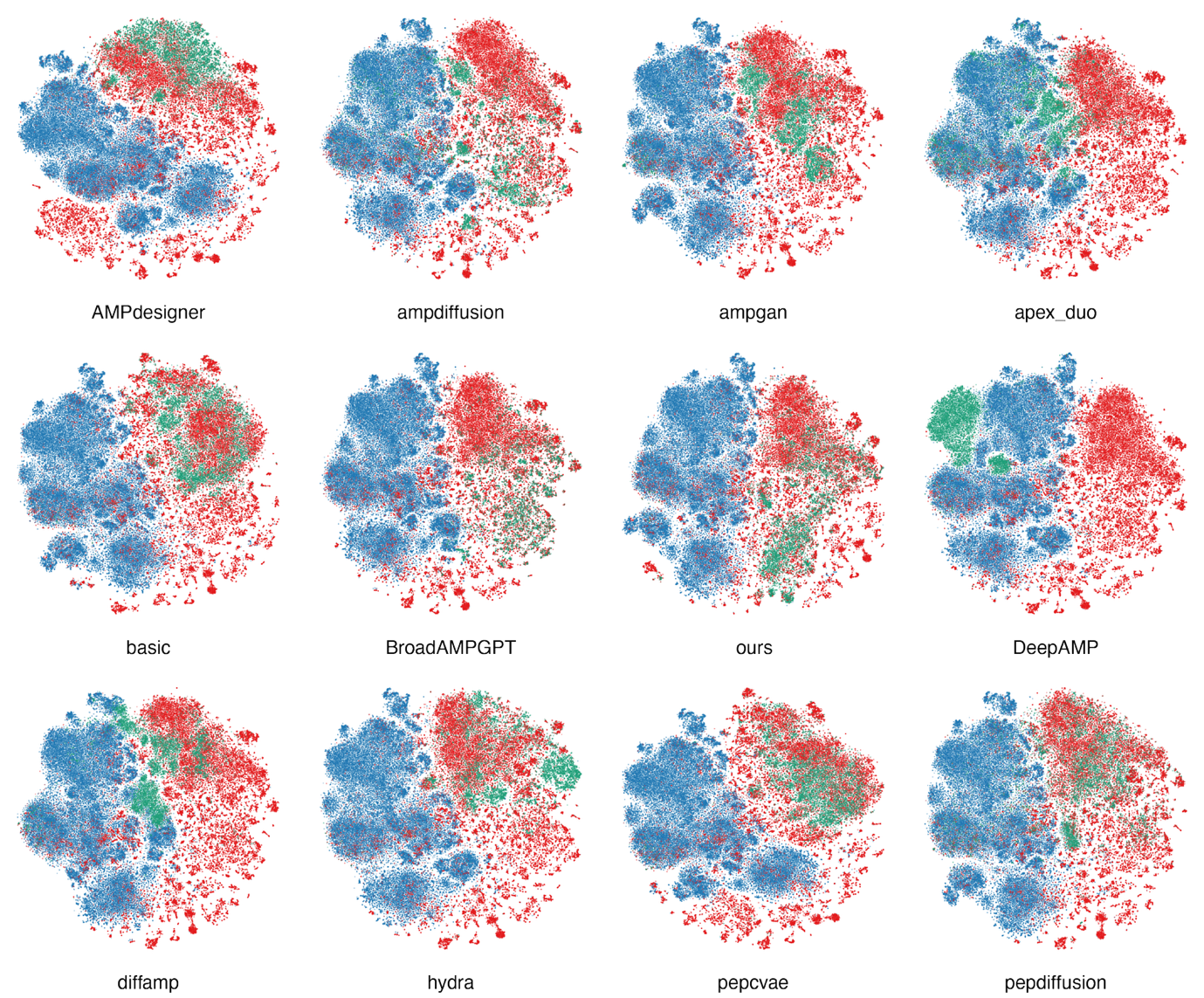

**Figure S11. t-SNE visualization of peptide sequence space (n = 10,000) from different generative models.** 10,000 peptides were randomly sampled from 26,000 peptides generated by our model and compared with an equal number from other methods, including AMPdesigner, ampdiffusion, AMP-GAN, ApexDuo, Basic, BroadAMPGPT, DeepAMP, DiffAMP, Hydra, PepCVAE, and PepDiffusion. Peptides were encoded using ESM-based embeddings, and t-SNE was used to project the high-dimensional representations into two-dimensional space, with colors indicating AMP-dissimilar peptides (blue), AMPs (red), and generated peptides (green).

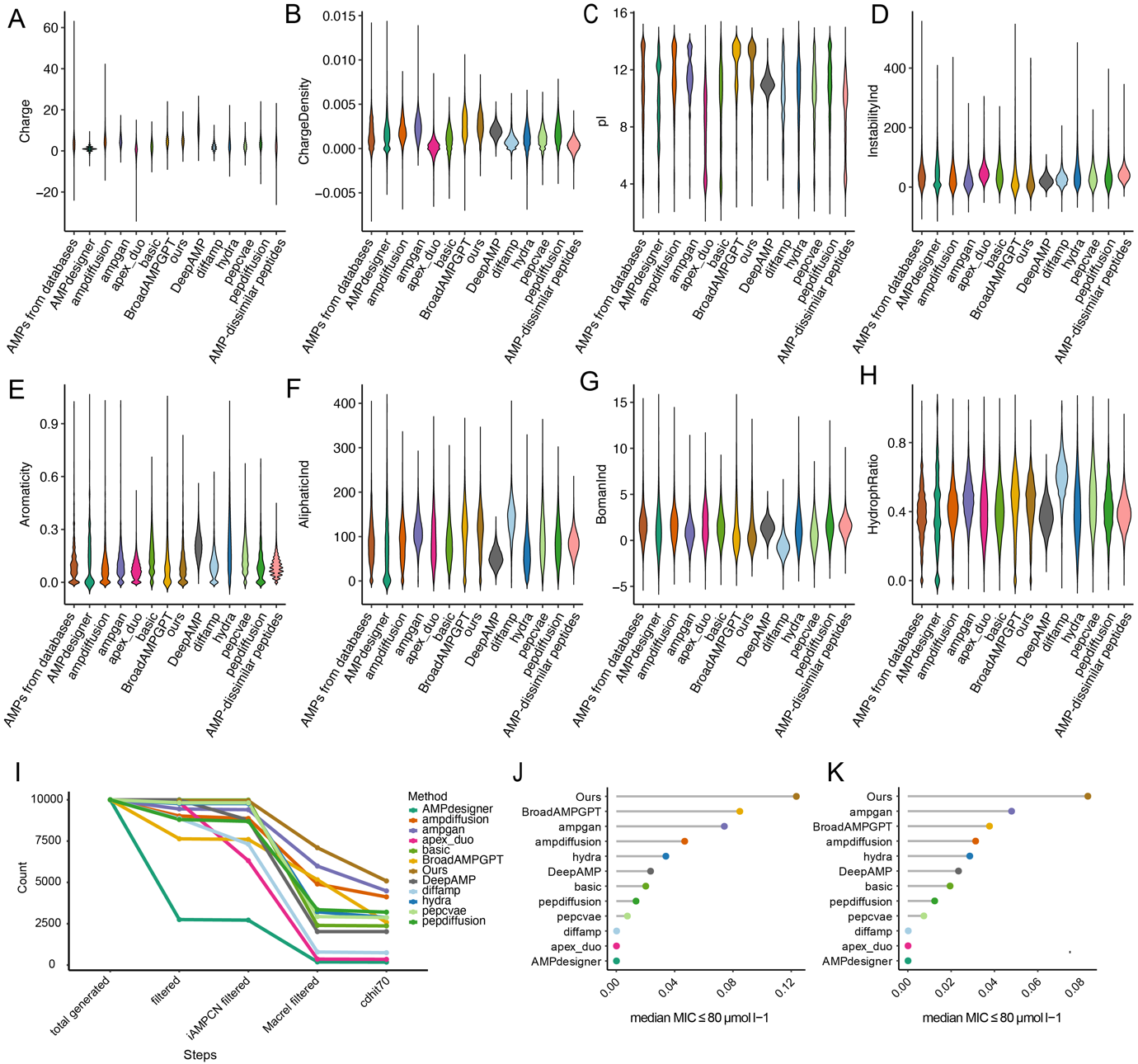

**Figure S12. Benchmarking of AMP generation methods across multiple physicochemical and filtering criteria. (A–H)** Violin plots showing the distributions of predicted properties of peptides generated by different methods, including **(A)** net charge, **(B)** charge density, **(C)** isoelectric point, **(D)** instability index (calculated based on dipeptide composition using empirically derived instability weights), **(E)** aromaticity [defined as the fraction of aromatic residues, phenylalanine (F), tryptophan (W), and tyrosine (Y) over other residues of the sequence], **(F)** aliphatic index, **(G)** Boman index, and **(H)** hydrophobic ratio **[defined as the proportion of alanine (A), cysteine (C), phenylalanine (F), isoleucine (I), leucine (L), methionine (M), and valine (V) residues]**. These physicochemical properties were calculated using the Global Analysis function in the modlamp Python package. Each colour corresponds to a different generative framework. **(I)** Success rate (fraction retained) for each method along a sequential filtering pipeline: (1) remove sequences containing the non-canonical/ambiguous residue “X” or with length <4; (2) apply iAMPCN to predict AMP likelihood and retain positives; (3) re-predict the retained set with Macrel and keep predicted AMPs; (4) filter out sequences with high similarity to known AMPs using CD-HIT at 70% sequence identity against the reference AMP database. Curves show the proportion of candidates remaining after each step for each model. **(J–K)** Success rates of AMP candidates under two alternative filtering pipelines. In both pipelines, candidate sequences were first evaluated using the APEX model, and peptides with an average median predicted MIC <80 μmol L^-1^ were retained. The success rate was defined as the percentage of retained peptides relative to the total number of generated sequences. Panel **(J)** shows the overall percentages without redundancy reduction, while panel **(K)** reports the corresponding values after clustering with CD-HIT at 70% sequence identity to remove peptides highly similar to known AMPs.
